# An amiRNA screen uncovers redundant CBF & ERF34/35 transcription factors that differentially regulate arsenite and cadmium responses

**DOI:** 10.1101/2020.12.30.424898

**Authors:** Qingqing Xie, Qi Yu, Timothy O. Jobe, Allis Pham, Chennan Ge, Qianqian Guo, Jianxiu Liu, Honghong Liu, Huijie Zhang, Yunde Zhao, Shaowu Xue, Felix Hauser, Julian I. Schroeder

## Abstract

Arsenic stress causes rapid transcriptional responses in plants. However, transcriptional regulators of arsenic-induced gene expression in plants remain less well known. To date, forward genetic screens have proven limited for dissecting arsenic response mechanisms. We hypothesized that this may be due to the extensive genetic redundancy present in plant genomes. To overcome this limitation, we pursued a forward genetics screen for arsenite tolerance using a randomized library of plants expressing >2,000 artificial microRNAs (amiRNAs). This library was designed to knock-down diverse combinations of homologous gene family members within sub-clades of transcription factor and transporter gene families. We identified six transformant lines showing an altered response to arsenite in root growth assays. Further characterization of an amiRNA line targeting closely homologous CBF and ERF transcription factors show that the CBF1,2 and 3 transcription factors negatively regulate arsenite sensitivity. Furthermore, the ERF34 and ERF35 transcription factors are required for cadmium resistance. Generation of CRISPR lines, higher-order T-DNA mutants, and gene expression analyses, further support our findings. These ERF transcription factors differentially regulate arsenite sensitivity and cadmium tolerance.

## Introduction

Arsenic (As) is a carcinogenic metalloid found widely in the environment (Clemens & Ma, 2016; Cooper et al., 2020). Arsenic toxicity decreases crop yields, and accumulation of arsenic in edible crop tissues can cause contamination of the food chain and lead to severe human health problems (Clemens, 2019; Rai, Lee, Zhang, Tsang, & Kim, 2019;Abedi & Mojiri, 2020; Palma-Lara et al., 2020). For example, rice grain can accumulate up to 2.24 mg/kg As and has become a dominant dietary exposure route to arsenic (As) (Gilbert-Diamond et al., 2011; Pan, Wu, Xue, & Hartley, 2014). Thus, understanding the molecular mechanisms of As uptake and accumulation in plants is an urgent priority for food security worldwide.

Recent studies have identified genes involved in the uptake, transport, and sequestration of arsenic in *Arabidopsis,* rice, and other plant species (Mendoza-Cozatl,Jobe, Hauser, & Schroeder, 2011; Abbas et al., 2018; Kumari, Rastogi, Shukla,Srivastava, & Yadav, 2018; Garbinski, Rosen, & Chen, 2019; Shri et al., 2019; Abedi & Mojiri, 2020). These include nutrient transporters that transport arsenite (As(III)) across plant membranes and can function in arsenic accumulation, including Nodulin 26-like Intrinsic Proteins (NIPs), Plasma membrane Intrinsic Proteins (PIPs), Natural Resistance-Associated Macrophage Protein OsNRAMP1, and Tonoplast Intrinsic Proteins (TIP) (Bienert et al., 2008; Isayenkov & Maathuis, 2008; Kamiya & Fujiwara, 2009; Kamiya et al., 2009; Mosa et al., 2012; Tiwari et al., 2014; Xu et al., 2015; Yang et al., 2015; Duanet al., 2016b; Lindsay & Maathuis, 2016). Furthermore, phosphate transporters (PHTs) play a role in arsenate (As(V)) uptake (Shin, Shin, Dewbre, & Harrison, 2004; Catarecha et al., 2007; Remy et al., 2012; LeBlanc, McKinney, Meagher, & Smith, 2013; Fontenot et al., 2015), and arsenate reductase enzymes are responsible for the reduction of As(V) to arsenite As(III) (Chao et al., 2014; Shi et al., 2016; Xu et al., 2017). The enzymes *γ*-glutamylcysteine synthetase (g-ECS), glutathione synthetase (GS), and phytochelatin synthase (PCS) are critical enzymes for As detoxification (Schmoger, Oven, & Grill, 2000;Li et al., 2004; Picault et al., 2006; Gasic & Korban, 2007; Guo, Dai, Xu, & Ma, 2008;Herschbach et al., 2010; Wojas, Clemens, Sklodowska, & Maria Antosiewicz, 2010;Hayashi et al., 2017). Additionally, the ATP Binding Cassette (ABC) transporters ABCC1 and ABCC2 transport As-GSH and As-PC complexes into vacuoles as a mechanism for arsenic accumulation (Song et al., 2010; Song et al., 2014; Hayashi et al., 2017). The inositol transporters (AtINT2 and AtINT4) in *A. thaliana* and OsNIP6;1 and OsNIP7;1 in rice play a role in arsenic transport into seeds (Duan et al., 2016a; Lindsay & Maathuis,2017).

Moreover, several regulatory proteins involved in plant responses to arsenic have been identified. For example, glutaredoxins (Grx) regulate As(V) reduction (Verma et al.,2017), and the transcription factors *WRKY6, WRKY45,* and *OsARM1* (Arsenite-Responsive Myb1) have been implicated in the regulation of arsenic transporters (Castrillo et al., 2013; Wang et al., 2014; Wang et al., 2017). The calcium-dependent protein kinase *(CPK31)* was reported to regulate *AtNIP1;1* (Ji et al., 2017), and *miR528* was found to be necessary for As(III) responses (Liu et al., 2015). Despite the above list of genes identified as being involved in arsenic responses in plants, many genes and mechanisms have yet to be determined.

Arsenic is known to cause large transcriptional responses in plants (Chakrabarty et al., 2009; Jobe et al., 2012; Castrillo et al., 2013; Srivastava, Srivastava, Sablok,Deshpande, & Suprasanna, 2015; Zvobgo et al., 2018; Huang et al., 2019). Research has suggested that transcriptional activators and transcriptional repressors play essential roles in arsenic-induced gene expression (Jobe et al., 2012; Castrillo et al., 2013; Wang et al., 2014; Wang et al., 2017). While WRKY transcription factors have been shown to regulate phosphate transporters that contribute to arsenic tolerance (Castrillo et al., 2013;Wang et al., 2014; Wang et al., 2017), additional transcription factors that mediate arsenic-induced gene expression and repressors remain unknown.

Forward genetic screens are powerful tools for identifying new genes. However, forward genetic screens have limitations due to the extensive over-lapping functions of closely related genes (“redundancy”) mediated by large gene families found in plant genomes (Arabidopsis Genome, 2000). Phenotypes linked to a single gene loss-of-function mutation have been found for less than 15% of the genes encoded in the *Arabidopsis* genome (Lloyd & Meinke, 2012; Cusack et al., 2020). A genome-wide analysis showed that approximately 75% of *Arabidopsis* genes are members of gene families (Hauser et al., 2013). Thus, partially overlapping gene functions, referred to as functional redundancies, have hampered the identification of new gene functions in forward genetic screens. In previous research, we computationally designed artificial microRNAs (amiRNAs) (Schwab, Ossowski, Riester, Warthmann, & Weigel, 2006), and synthesized 22,000 amiRNAs designed to combinatorially co-silence closely homologous gene clade members (Hauser et al., 2013). These amiRNAs were separated into 10 amiRNA libraries for genome-wide knockdown of homologous gene family members based on the predicted functional classes of the genes they target (Hauser et al., 2013;Hauser et al., 2019). Using these 22,000 amiRNAs, libraries were synthesized (Hauser et al., 2013) and used to transform *Arabidopsis*. To date, 14,000 T2 generation lines have been generated (Hauser et al., 2019). Recent studies have used this resource to discover new genes and determine their functions in plant hormone signaling and transport, illustrating the power of this system (Hauser et al., 2013; Zhang et al., 2018; Hauser et al., 2019; Takahashi et al., 2020).

To overcome the genetic redundancy limitations, we screened amiRNA seed libraries targeting two different gene classes (1: DNA and RNA binding proteins and 2: transporters and channels) for arsenic tolerance. Here, we report the identification of new transcription factors and transporters that play crucial roles in arsenic responses.

## Material and Methods

### Genetic screen for mutants with altered arsenite response

Wild type (Col-0) and individually isolated T2 lines from the amiRNA libraries (Hauser et al., 2013) were surface sterilized, stratified at 4°C for 48 hrs in the dark, and germinated under a 16 hrs light/ 8 hrs dark photoperiod.

For root length assays(Lee, Chen, & Schroeder, 2003; Jobe et al., 2012; Mendoza-Cozatl et al., 2014), 1/2 MS plates (1/2 MS, 0.5 g/L MES, 0.5% sucrose, 1% phytagel, adjusted with KOH to pH 5.5) were supplemented with/without 10 μM sodium arsenite and allowed to grow vertically for 14 days. The root lengths of seedlings were measured using Image J. The same protocol was performed on the next generation of (T3) seeds for candidate lines showing a phenotype.

Root length assays for *erf34×35, CRISPRcbf1/2/3* were performed on minimal medium (5 mM KNO3, 2.5 mM H3PO4, 1 mM Ca(NO3)2, 2 mM MgSO4, 1 mM MES, adjusted with KOH to pH 5.5, 1% phytagel) without microelements supplied with and without 10 μM arsenite.

### Identification of amiRNA sequences

Genomic DNA for candidate amiRNA lines were extracted(Kang, Cho, Yoon, & Eun,1998) and used as templates, and the primers (primers pha2804f and pha3479r(Hauser et al., 2019), Supplemental Table 1) were used to amplify the PCR product. The purified PCR samples were sent for sequencing to identify the amiRNA present(Hauser et al.,2019). The genes targeted by the amiRNA could be putatively identified by using the “Target Search Function” on the WMD3 website (Ossowski, Fitz, Schwab, Riester, & Weigel).

### Plasmid Construction

To test whether the phenotype of the amiRNA lines was caused by the knockdown of target genes, and to confirm that the phenotypes were not caused by the position of the amiRNA insertion, we generated retransformed lines using the genomic DNA of candidate amiRNA lines as a template. We named these lines Re-10-9. The primers RE-AMI-F and RE-AMI-R were used to amplify the original amiRNA and inserted it into pDONR221^®^ using BP Clonase II® and then recombined into pFH0032 (Hauser et al.,2013; Hauser et al., 2019) using LR Clonase II® (Invitrogen, Carlsbad,CA, USA). (Supplemental Table 1).

We generated *erf34xerf35* by crossing the T-DNA lines. Primers for genotyping were designed using the website (http://signal.salk.edu/tdnaprimers.2.html). A list of primers used in this study is provided in Supplemental Table 2.

We used CRISPR/Cas9 gene editing technology (Gao & Zhao, 2014; Gao et al.,2015; Gao, Chen, Dai, Zhang, & Zhao, 2016) to generate *cbf1/2/3* knockout mutants. Guide RNA 1 and 2 were used to delete the CBF1/2/3 genes, which are tandemly arrayed on chromosome IV. The target sequence for target 1 was CCGATTACGAGCCTCAAGGCGG, and the sequence for target 2 was CCGGAACAGAGCCAAGATGCGT (PAM sites are underlined). The GT1-3 primers were used to isolate the homozygous lines for *CRISPR cbf1/2/3.* GT1 is the forward primer, GT2 and GT3 are reverse primers. The homozygous plants were isolated as: GT1+GT2 has band, and no band for GT1+GT3 (Supplemental Figure 3A and B).

### Plant Transformation

Sequenced plasmids were transformed into *Agrobacterium tumefacians* strain GV3101 and pSoup was used as the helper plasmid for pFH0032. *Arabidopsis thaliana* was transformed as previously described using the floral dip method(Clough & Bent,1998).

### Total RNA isolation and quantitative RT-PCR

For RT-qPCR, plants were grown vertically on 1/2 MS media for 14 days and transfer to liquid treatment (1/2 MS, 0.5 g/L MES, 0.5% sucrose, pH 5.5-5.8 with/without 10 μM Arsenite) for 3 days. Leaves and roots were separated, and RNA was prepared using the Spectrum Plant Total RNA Kits (Sigma-Aldrich). cDNA was then prepared from RNA using Maxima H Minus First Strand cDNA Synthesis Kit (Thermo Scientific™) for RT-PCRs. Transcript abundance was then determined with SYBR® Green (Bio-Rad, MO, USA) using a Bio-Rad CFX96 Real-Time System with the following conditions: 95 °C for 5 min, then 50 cycles of 95°C for 15 s, 52°C for 15 s, 72°C for 1 min, then 72°C for 5 min. Primers used for qPCR(Versaw & Harrison, 2002; B. Guo et al., 2008; Remy et al., 2012;Lapis-Gaza, Jost, & Finnegan, 2014) are listed in Supplemental Table 3.

## Results

### Genetic screen for mutants with altered arsenite response

In previous research, arsenite was found to decrease primary root growth, inhibit seedling development, and cause chlorosis (Liu, Zhang, Shan, & Zhu, 2005; Yoon, Lee, & An, 2015; Qian et al., 2018). We tested a range of arsenite concentrations on 1/2 MS media-containing plates to establish the concentration at which wild-type seedling root growth was decreased by 40%-60% compared to control plates. We chose a concentration of 10 μM arsenite for root length screening.

In total, ~2,000 T2 amiRNA expressing *Arabidopsis* transformant lines were screened. These amiRNA lines were screened individually in root growth assays. Seed transformed with amiRNA libraries targeting homologous genes that encode DNA and RNA binding proteins and transporter family proteins were screened(Hauser et al., 2013;Hauser et al., 2019). In the primary screen, 15 seeds per line were screened for differences in root growth compared to wild-type plants grown in parallel. Candidate lines showing an altered arsenite response were validated using the same phenotyping method in the next generation (T3). Isolated amiRNA lines with reproducible phenotypes were then analyzed for their putative target genes by sequencing and functional characterization. After screening these 2,000 amiRNA lines, six candidate amiRNA lines were identified with robust phenotypes. Their target genes and gene definitions are shown in Table 1 (e.g. Figure 1A-C and Supplemental Figure 1). In the present study, two amiRNA lines, amiRNA *4-138* and ami *10-9* were chosen for further characterization.

**Table 1.**
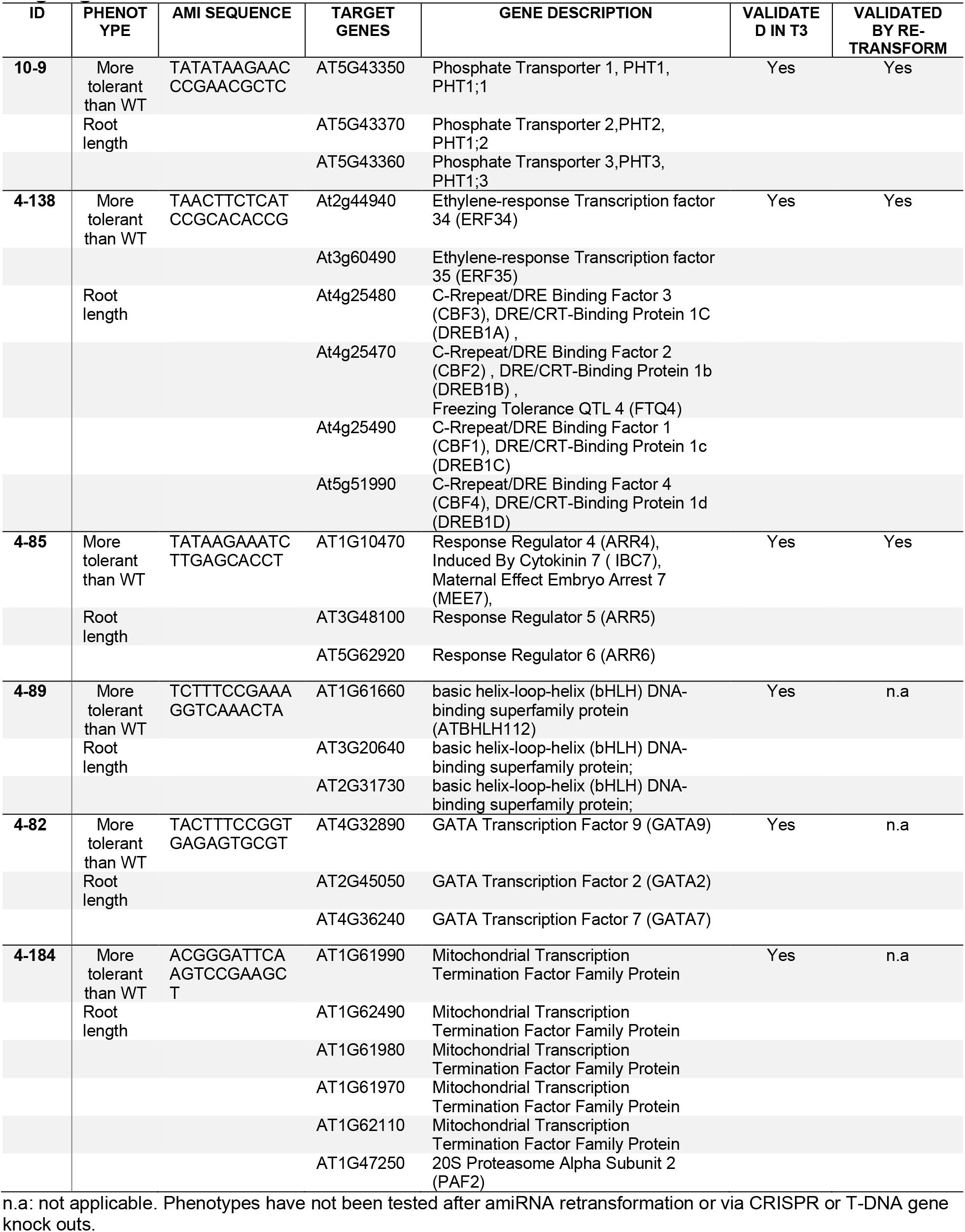
Candidate amiRNA lines with altered response to Arsenite and their target genes

**Figure 1.**
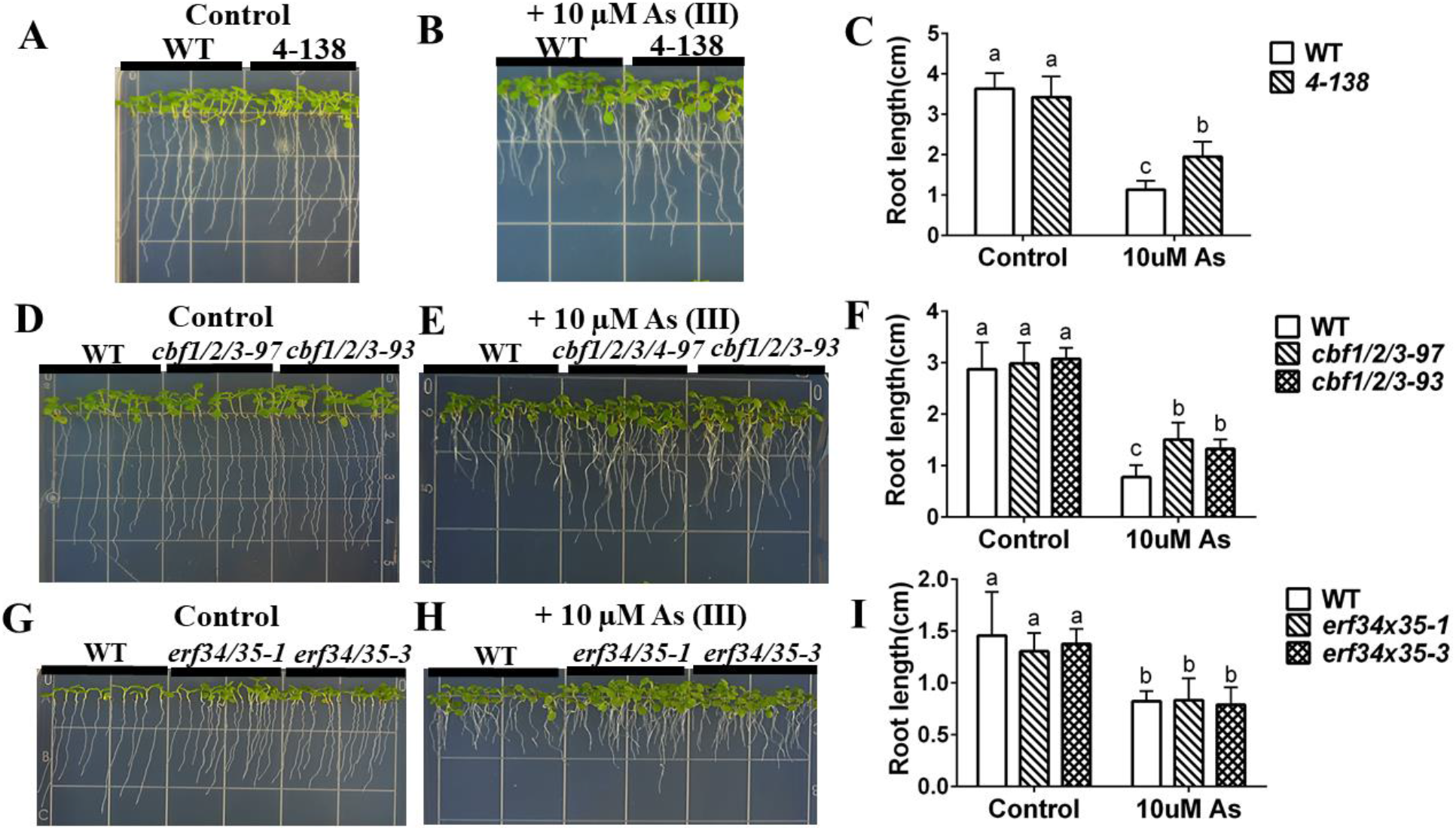
amiRNA *4-138* targets ERF and CBF transcription factors, shows arsenite resistance, and phenocopies CRISPR lines targeting *CBF1,2,3* genes but not *erf34/35* double T-DNA mutants. (A), (B) amiRNA *4-138* is more tolerant than WT to arsenite treatment: WT and amiRNA *4-138* lines were grown vertically on 1/2 MS with 0.5% sucrose plates without (control) or with 10 μM arsenite for 10 days; (C) Primary root lengths of amiRNA *4-138* seedlings were significantly longer than WT on arsenite-containing plates. (n=11 seedlings, means ± s.e.m.). (D), (E) CRISPR lines targeting *CBF1/2/3* genes show arsenite resistance of root growth: CRISPR lines 97 and 93 targeting *CBF1/2/3* were grown vertically on minimal media plates with 1% sucrose plates without (control) or with 10 μM arsenite for 7 days; (F) Primary root lengths of amiRNA *4-138* were significantly longer than WT on arsenite-containing plates. (n=11 seedlings, means ± s.e.m.). (H), (I) T-DNA double mutant in *ERF34* and *ERF35* genes showed similar root length to wild type under both control and arsenite treatment: WT and double mutant of *erf34/35* line 1 and 3 were grown vertically on minimal plates with 1% sucrose plates without (control) or with 10 μM arsenite for 7 days. (I) Averaged primary root length (n=16 seedlings, means ± s.e.m.). Letters at the top of columns are grouped based on two-way ANOVA and Tukey’s multiple comparisons test, P < 0.05).

### amiRNA line *4-138* targets ERF and CBF transcription factors and is tolerant to arsenite

A mutant was isolated in the amiRNA screen showing an increased resistance to arsenite in root growth assays. Primary root lengths of amiRNA *4-138* seedlings were not different from WT on control plates but were significantly longer than WT on arsenite-containing plates (Figure 1A-C).

The isolated amiRNA *4-138* is predicted to target two different *ERF* transcription factor sub-families, specifically *ERF 34, 35, and CBF 1, 2, 3, & 4.* To determine which of these putative targets contributes to the arsenite-tolerant phenotype, we first screened single mutants and observed only weak or no phenotypes (see Supplemental figure 2A and C). Then double mutants were generated using T-DNA lines *erf34-1* (SALK_020979C) and *erf35-1* (SALK_111486C) (see Supplemental figure 4A and B). Furthermore, CRIPSR lines were created for *cbf 1/2/3.* Two independent *cbf 1/2/3* CRISPR deletion alleles were isolated. The CRISPR lines targeting *CBF1/2/3* did not show any visible phenotype under control conditions in plate growth assays (Figure 1D). However, both *cbf1/2/3* CRISPR alleles showed arsenite resistance in seedling root growth assays (Figure 1D-F). In contrast, *erf34/35* double mutants exhibited arsenite sensitivity similar to WT (Figure 1G-I). These data suggest that the arsenite-tolerant phenotype of amiRNA line *4-138* is caused by the knockdown of *CBF 1, 2, 3* transcription factors, rather than the *ERF 34* and *35* transcription factors.

### CBF transcription factors and PHT transporters are down-regulated in response to arsenic treatment and are misregulated in the *cbf1,2,3* mutant

The enhanced arsenic resistance of the *cbf1/2/3* mutant alleles indicates that CBF transcription factors may function as negative regulators of arsenic resistance. To investigate whether the transcript levels of *CBF1, CBF2, and CBF3* were affected by arsenite or arsenate stress, we measured the transcript levels of these transcription factors using qPCR. Interestingly, the transcript levels of these three genes decreased in seedlings exposed after 10 days of growth to 10 μM As(III) or 250 μM As (V) for 3 days (Figure 2A-C).

**Figure 2.**
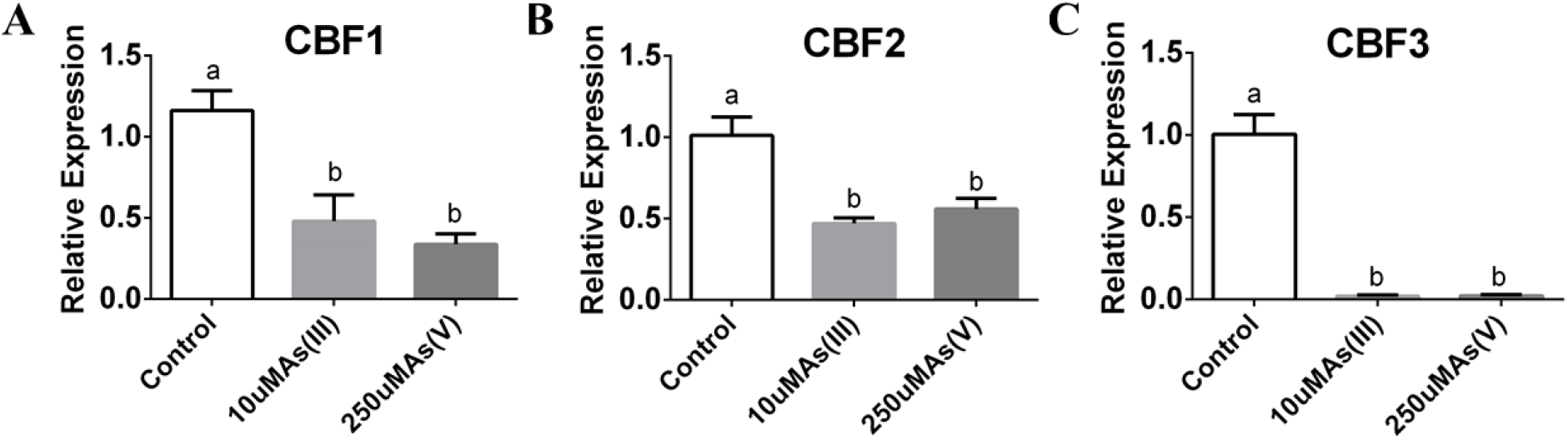
*CBF1, CBF2* and *CBF3* are transcriptionally downregulated by arsenite and arenate in the roots. (A-C) Transcript levels of *CBF1, 2, 3* under arenite and arsenate treatment are significantly lower than control condition in roots of WT: WT plants were grown on 1/2 MS for 10 days and transferred to 1/2 MS plates with or without 10 μM sodium arsenite or 250 μM sodium arsenate for 3 days. Root tissues were harvested separately, followed by RNA extraction and cDNA synthesis, and the relative expression levels were detected by qRT-PCR. (n=3 samples, means ± s.e.m.). Letters at the top of columns are grouped based on two-way ANOVA and Tukey’s multiple comparisons test, P < 0.05).

We examined possible DNA-binding targets of the CBF transcription factors. The Plant Cistrome Database (O’Malley et al., 2016) has used DNA affinity purification sequencing (DAP-seq) for resolving possible motifs and peaks for DNA binding domains of 529 recombinant *Arabidopsis* transcription factors. A list of genes putatively targeted by the CBF1-4 transcription factors was generated (O’Malley et al., 2016) (Supplemental data 2). Interestingly CBF1-4 binding was observed in the promoters of several phosphate transporters (PHT), including *PHT1;1, PHT1;3, PHT1;6, PHT1;7, PHT1;8, PHT1;9, PHT3;3, PHT4;2, PHT2;1 PHT4;3* in these DAP-seq experiments (Table 2). Thus, we investigated whether these transporter genes might be targets of CBF1-4. Additionally, *PHT1;1, PHT1;2* and *PHT 1;3* were the target genes of the ami *10-9* line that was isolated in our forward genetics screen, which unexpectedly showed an arsenite tolerant phenotype (Table 1), as described later.

**Table 2.** Target phosphate transporters of CBF1, 2, 3 and 4

|  | TARGET GENES | GENE SYMBOL | GENE DESCRIPTION |
| --- | --- | --- | --- |
| **CBF1** | AT5G43340 | PHT1;6 | Phosphate Transporter 1;6 |
|  | AT5G43360 | PHT1;3 | Phosphate Transporter 1;3 |
|  | AT1G20860 | PHT1;8 | Phosphate Transporter 1;8 |
|  | AT1G76430 | PHT1;9 | phosphate transporter 1;9 |
|  | AT2G17270 | PHT3;3 | phosphate transporter 3;3 |
|  | AT2G38060 | PHT4;2 | phosphate transporter 4;2 |
|  | AT3G26570 | PHT2;1 | phosphate transporter 2;1 |
|  | AT3G46980 | PHT4;3 | phosphate transporter 4;3 |
| **CBF2** | AT1G76430 | PHT1;9 | phosphate transporter 1;9 |
|  | AT2G38060 | PHT4;2 | phosphate transporter 4;2 |
|  | AT3G26570 | PHT2;1 | phosphate transporter 2;1 |
|  | AT3G46980 | PHT4;3 | phosphate transporter 4;3 |
|  | AT5G43360 | PHT1;3 | Phosphate Transporter 1;3 |
| **CBF3** | AT1G76430 | PHT1;9 | phosphate transporter 1;9 |
|  | AT3G46980 | PHT4;3 | phosphate transporter 4;3 |
|  | AT3G26570 | PHT2;1 | phosphate transporter 2;1 |
| **CBF4** | AT1G20860 | PHT1;8 | Phosphate Transporter 1;8 |
|  | AT1G76430 | PHT1;9 | phosphate transporter 1;9 |
|  | AT2G38060 | PHT4;2 | phosphate transporter 4;2 |
|  | AT3G26570 | PHT2;1 | phosphate transporter 2;1 |
|  | AT3G46980 | PHT4;3 | phosphate transporter 4;3 |
|  | AT3G54700 | PHT1;7 | phosphate transporter 1;7 |
|  | AT5G43350 | PHT1;1 | phosphate transporter 1;1 |
|  | AT5G43360 | PHT1;3 | Phosphate Transporter 1;3 |

To determine whether transcript levels of PHT transporters are affected in the CRISPR *cbf1/2/3* mutants, we measured the expression levels of the *PHT* transporter mRNAs. We found that the transcript levels of *PHT1;1,1;2, 1;3* decreased significantly in wild-type Col-0 (WT) roots in response to arsenite treatment (Figure 3A-C). Furthermore, *PHT 1;1, 1;2,* and *1;3* transcript levels in the CRISPR *cbf1/2/3* mutants were much lower in the roots under control conditions (Figure 3A-C). However, no differences between WT and CRISPR *cbf1/2/3* mutants under arsenite treatment were observed, apparently because the transcripts were already expressed at low levels in arsenic-free controls (Figure 3A-C).

**Figure 3.**
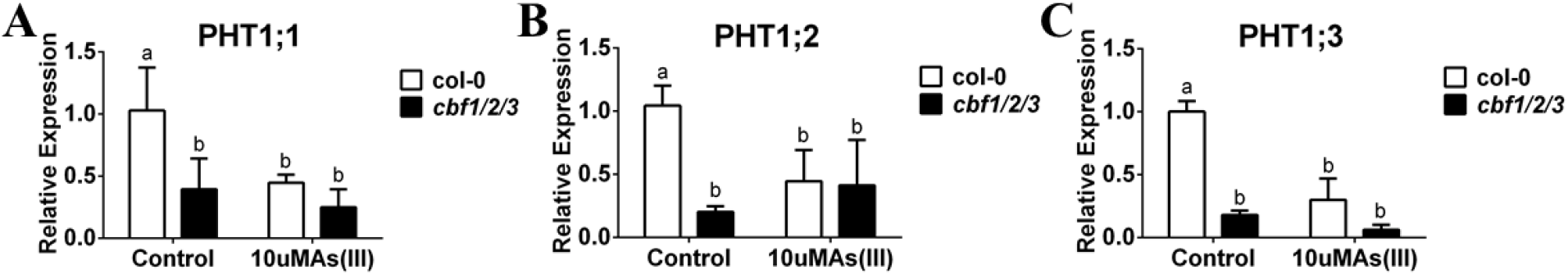
*PHT transporter* transcript levels are down-regulated in response to arsenite treatment and misregulated in roots of CRISPR *cbf 1,2,3* mutant. (A-C) *PHT1;1; PHT1;2 and PHT1;3* are reduced significantly in roots by arsenite treatment, and *PHT 1;1; PHT1;2 and PHT1;3* transcript levels in CRISPR *cbf1/2/3* mutants are much lower in the roots under control conditions: WT and CRISPR *cbf1/2/3* mutants were grown on 1/2 MS for 10 days and transferred to 1/2 MS plates with or without 10 μM arsenite for 3 days. Root tissues were harvested separately, followed by RNA extraction and cDNA synthesis, and the relative expression levels were detected by qRT-PCR. (n=3 samples, means ± s.e.m.). Letters at the top of columns are grouped based on two-way ANOVA and Tukey’s multiple comparisons test, P < 0.05).

Because PHT transporters are known to transport phosphate and arsenate (Shin et al., 2004; Catarecha et al., 2007; Remy et al., 2012; LeBlanc et al., 2013; Fontenot et al.,2015), the transcript levels of *PHTs* were also measured under arsenate treatment. *PHT 1;1, 1;2, 1;3* transcript levels were significantly decreased in wild-type roots in response to arsenate treatment (Figure 4A-C). In *cbf1/2/3* triple mutants, the control non-stress transcript levels of these *PHTs* were decreased. These data suggest that CBF transcription factors play a role in upregulating several PHT transcripts under non-stress conditions.

**Figure 4.**
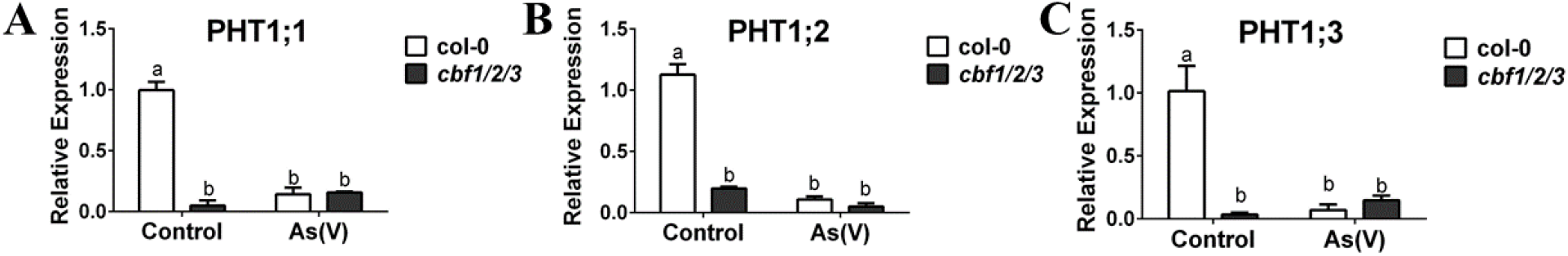
*PHT transporter* transcript levels are down-regulated in response to arsenate treatment and misregulated in roots of CRISPR *cbf 1,2,3* mutant. (A-C) *PHT1;1; PHT1;2, PHT1;3* are reduced significantly in roots by arsenate treatment and *PHT* transcript levels in CRISPR *cbf1/2/3* mutants are much lower in the roots under control conditions: WT and CRISPR *cbf1/2/3* mutants were grown on 1/2 MS for 10 days and transferred to 1/2 MS plates with or without 250 μM sodium arsenite for 3 days. Root tissues were harvested separately, followed by RNA extraction and cDNA synthesis and the relative expression levels were detected by qRT-PCR. (n=3 samples, means ± s.e.m.). Letters at the top of columns are grouped based on two-way ANOVA and Tukey’s multiple comparisons test, P < 0.05).

### amiRNA 4-138 is sensitive to cadmium

Because arsenic and the toxic heavy metal cadmium elicit similar transcriptional responses (Abercrombie et al., 2008; Jobe et al., 2012; Shukla et al., 2018), we investigated whether amiRNA *4-138* also exhibits a cadmium-dependent phenotype. Interestingly, in contrast to As exposure (Figure 1A-C), seedlings of the amiRNA *4-138* line were more sensitive to cadmium compared to WT controls (Figure 5A-C). To determine which of the five predicted amiRNA targets contributes to the cadmium sensitivity, we evaluated root growth of the CRISPR *cbf1/2/3* and *erf34/35* mutant alleles in response to cadmium exposure. The two independent CRISPR alleles targeting *CBF1/2/3* genes showed cadmium sensitive phenotypes that were only slightly more severe than WT controls (Figure 5D-F). Interestingly, however, the two *erf34/35* double mutant alleles exhibited strong cadmium sensitivities compared to WT controls (Figure 5G-H). These data indicate that the *ERF 34, 35* transcription factors and possibly in part the *CBF 1, 2, 3* contribute to the cadmium sensitivity of the *4-138* amiRNA line.

**Figure 5.**
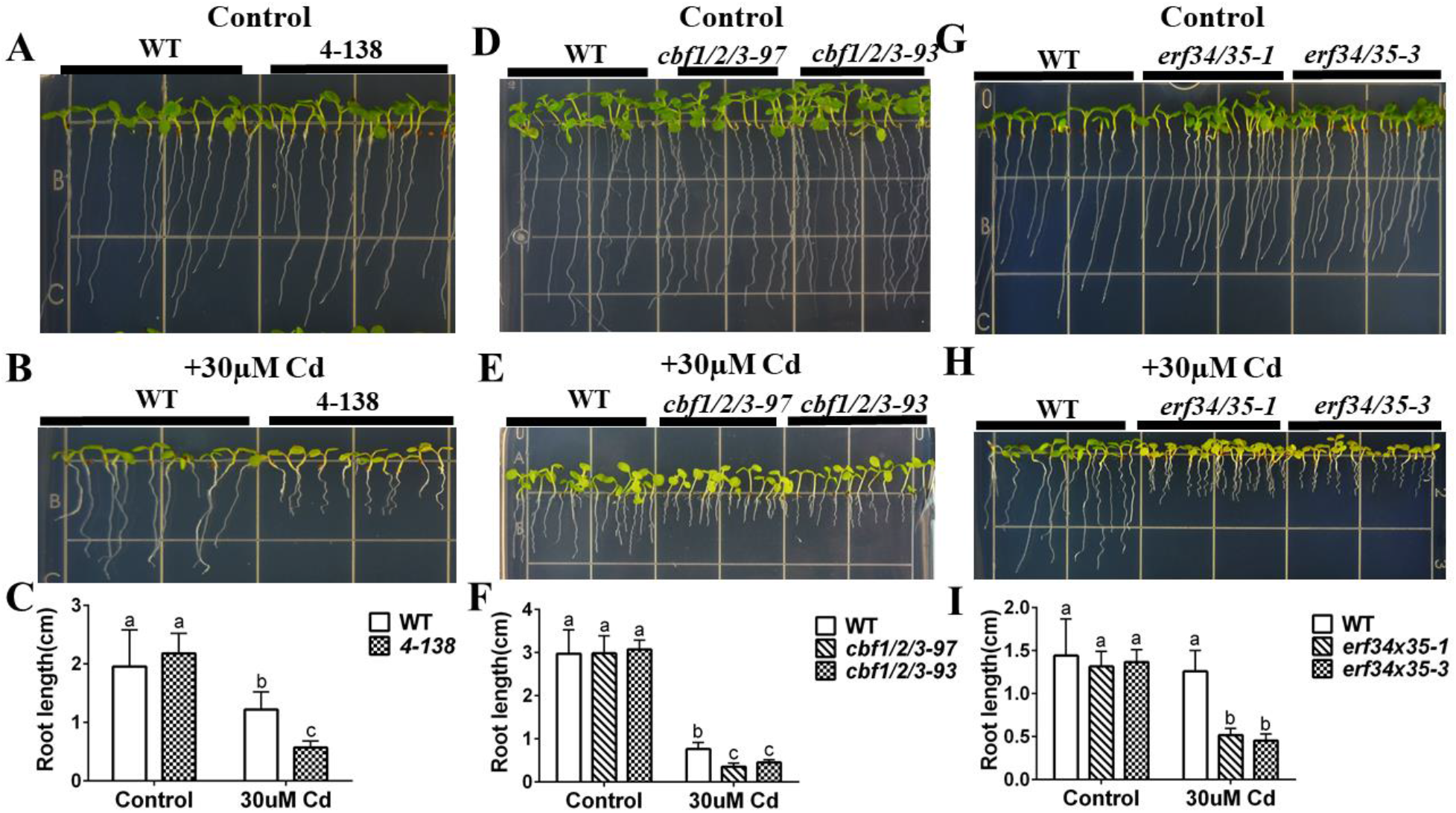
amiRNA *4-138* line and *erf34/erf35* double mutant are sensitive to cadmium. (A), (B) amiRNA line *4-138* shows enhanced cadmium sensitivity in root growth and yellowing of leaves compared to WT: WT and *4-138* amiRNA lines were grown vertically on minimal plates with 1% sucrose plates without (control) or with 30 μM cadmium for 7 days; (C) Primary root lengths of the amiRNA *4-138* line were significantly shorter than WT on cadmium-containing plates. (n=16 seedlings, means ± s.e.m.). (D), (E) CRISPR lines targeting *CBF1/2/3* genes show cadmium sensitive phenotypes. WT and CRISPR lines targeting *CBF1/2/3* genes were grown vertically on minimal plates with 1% sucrose plates without (control) or with 30 μM cadmium for 7 days; (F) Primary root length of CRISPR lines in *CBF1/2/3* genes compared to WT on control and cadmium-containing plates. (n=11 seedlings, means ± s.e.m.). (G), (H) *erf34/35* double mutant shows cadmium sensitive phenotype: WT and *erf34/35-3* T-DNA double mutant were grown vertically on minimal plates with 1% sucrose plates without (control) or with 30 μM cadmium for 7 days; (I) Primary root length of *erf34/35* compared to WT on control and cadmium-containing plates. (n=17 seedlings, means ± s.e.m.). Letters at the top of columns are grouped based on two-way ANOVA and Tukey’s multiple comparisons test, P < 0.05).

### Reduced *NRT 1.8* transcript levels in *erf34/35* in response to Cd

A list of candidate genes targeted by the *ERF34* and *ERF35* transcription factors were generated by searching The Plant Cistrome Database (O’Malley et al., 2016). The database was compared to genes significantly (p<0.05) induced more than 2-fold after 2 hours of exposure to 50 μM Cd^2+^ (Weber, Trampczynska, & Clemens, 2006). We examined possible DNA binding targets of ERF34 and ERF35, then analyzed which of these genes showed transcriptional regulation in response to cadmium exposure (Supplemental Figure 5). Based on the putative ERF34 and ERF35 targets from the Plant Cistrome Database, a list of candidate ERF34 and ERF35 targets also showing a transcriptional response to cadmium exposure was generated (Table 3). A previously identified nitrate transporter, NRT1.8, is a putative target of ERF34 and ERF35, based on analyses of the Plant Cistrome Database (O’Malley et al., 2016). Furthermore, the expression of *NTR1.8* was induced 29-fold by cadmium treatment in WT (Weber et al.,2006) (Table 3). NRT1.8 belongs to the nitrate transporter (NRT1) family and functions in nitrate removal from the xylem sap (Li et al., 2010). NRT1.8 affects the content of Cd in the xylem sap and mediates tolerance to cadmium (Li et al., 2010).

**Table 3.** Genes significantly (p<0.05) induced >2 folds after 2 hours of exposure to 50 μM Cd^2+^ in *A. thaliana* roots and targeted by EFR034 and ERF035.

| Gene ID | Gene Symbol | Gene description | Fold change | p-value |
| --- | --- | --- | --- | --- |
| AT4G21680 | NRT1.8 | NITRATE TRANSPORTER 1.8 | 29 | 0.0406 |
| AT4G14680 | APS3 | Pseudouridine synthase/archaeosine transglycosylase-like family protein | 12.3 | 0.0401 |
| AT1G62300 | WRKY6 | WRKY family transcription factor | 5.6 | 0.0327 |
| AT4G01950 | GPAT3 | glycerol-3-phosphate acyltransferase 3 | 4.3 | 0.0367 |
| AT4G17500 | ERF-1 | ethylene responsive element binding factor 1 | 3.9 | 0.0483 |
| AT3G12580 | HSP70 | heat shock protein 70 | 3.4 | 0.0327 |
| AT1G08920 | ESL1 | ERD (early response to dehydration) six-like 1 | 3.3 | 0.0483 |
| AT2G32560 | AT2G32560 | F-box family protein | 3 | 0.0483 |
| AT1G78820 | AT1G78820 | D-mannose binding lectin protein with Apple-like carbohydrate-binding domain-containing protein | 2.9 | 0.04 |
| AT3G25230 | ROF1 | rotamase FKBP 1 | 2.7 | 0.0401 |
| AT4G26080 | ABI1 | Protein phosphatase 2C family protein | 2.6 | 0.0483 |
| AT3G47960 | GTR1 | Major facilitator superfamily protein | 2.5 | 0.0483 |
| AT2G18690 | AT2G18690 | transmembrane protein | 2.4 | 0.05 |
| AT2G41800 | AT2G41800 | imidazolonepropionase (Protein of unknown function, DUF642) | 2.4 | 0.0483 |
| AT2G21130 | AT2G21130 | Cyclophilin-like peptidyl-prolyl cis-trans isomerase family protein | 2.3 | 0.0418 |
| AT2G41410 | AT2G41410 | Calcium-binding EF-hand family protein | 2.3 | 0.0483 |
| AT5G16600 | MYB43 | myb domain protein 43 | 2.3 | 0.0483 |
| AT3G46130 | MYB48 | myb domain protein 48 | 2.2 | 0.0485 |

We examined cadmium-dependent *NRT1.8* transcript levels to determine whether ERF34 and 35 affect *NRT1.8* expression. *NRT1.8* transcripts were detected at similar levels in both WT and the *erf34/35* double mutant seedlings under control conditions (Figure 6). *NRT1.8* was highly induced by Cd treatment in WT seedlings, consistent with previous findings (Weber et al., 2006; Li et al., 2010). However, cadmium did not significantly upregulate *NRT1.8* expression in *erf34/35* double mutant seedlings (Figure 6). The relatively low induction of *NRT1.8* may contribute to the Cd sensitive phenotype of the *erf34/35* double mutant.

**Figure 6.**
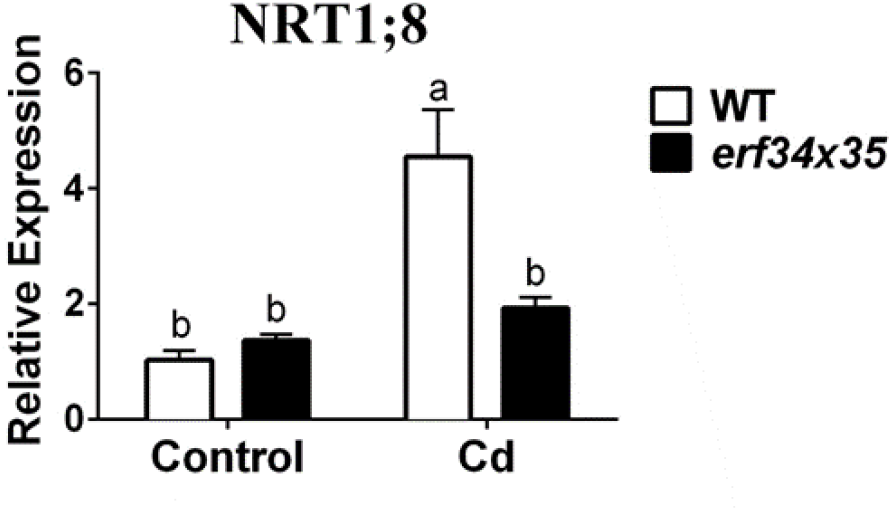
ERF34 and ERF35 are required for *NRT1;8* upregulation in response to cadmium. *NRT1;8* transcript up-regulation is impaired in roots of *erf34/35* double mutant: WT and *erf34/35* double mutants were subjected to the same treatment as in Figure 5 (G) and (H) (n=3; p<0.0001);

### ami10-9 is insensitive to arsenic and targets three phosphate transporters (PHTs)

In our forward genetics amiRNA screen for arsenic sensitivity, we also identified an amiRNA line *10-9* as showing arsenite resistance in root growth assays, that was similar to the amiRNA *4-138.* In the absence of arsenic, seedlings from this line have a similar primary root length as WT plants (Figure 7A). However, this line showed a reduced sensitivity compared to WT seedlings when grown on media containing 10 μM arsenite. Primary root lengths of amiRNA line *10-9* seedlings were almost twice as long as WT plants when grown on medium containing 10 μM As(III) (Figure 7B). A similar As insensitive phenotype of amiRNA line *10-9* was found on media containing arsenate (Figure 8A-C).

**Figure 7.**
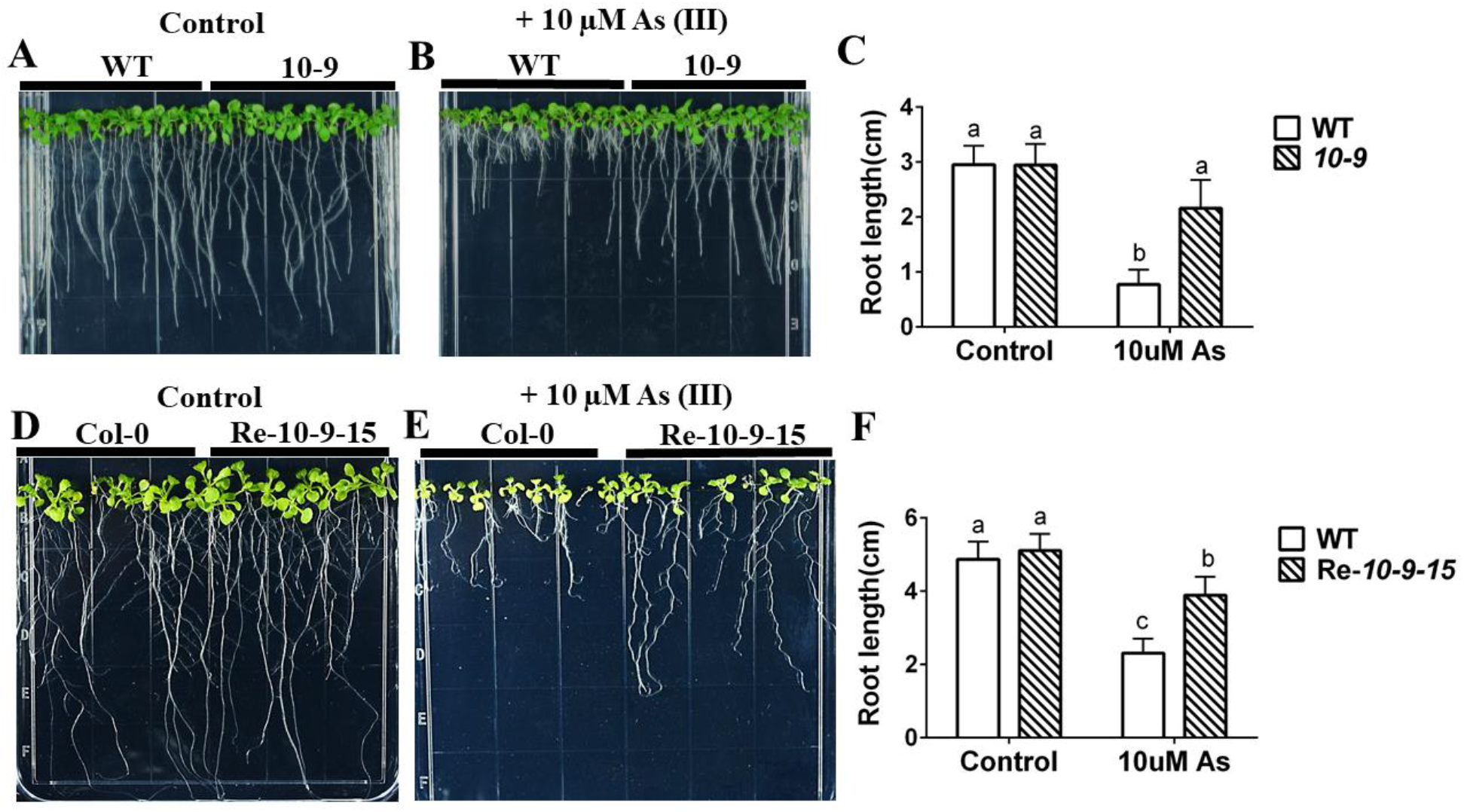
amiRNA *10-9* targets phosphate transporters and exhibits arsenite resistance. Arsenite resistance phenotype confirmed in amiRNA retransformation lines. (A), (B) Root growth of amiRNA *10-9* is more tolerant than WT to arsenite treatment: WT and amiRNA *10-9* lines were grown vertically on 1/2 MS with 0.5% sucrose plates without (control) or with 10 μM arsenite for 10 days; (C) Primary root lengths of WT and amiRNA *10-9* were significantly longer than WT on arsenite plates and no different on control plates. (n=15 seedlings, means ± s.e.m.). (D), (E) amiRNA retransformation lines with the same amiRNA as the amiRNA *10-9* line are more tolerant than WT in root growth assays: WT and amiRNA *10-9* retransformation line 15 were germinated directly on 1/2 MS with 0.5% sucrose plates without (control) or with 10 μM arsenite for 14 days; (F) Primary root lengths of retransformation amiRNA *Re-10-9-15* were significantly longer than WT on arsenite-containing plates and no clear difference was observed on control plates. (n=9 seedlings, means ± s.e.m.). Letters at the top of columns are grouped based on two-way ANOVA and Tukey’s multiple comparisons test, P < 0.05).

**Figure 8.**
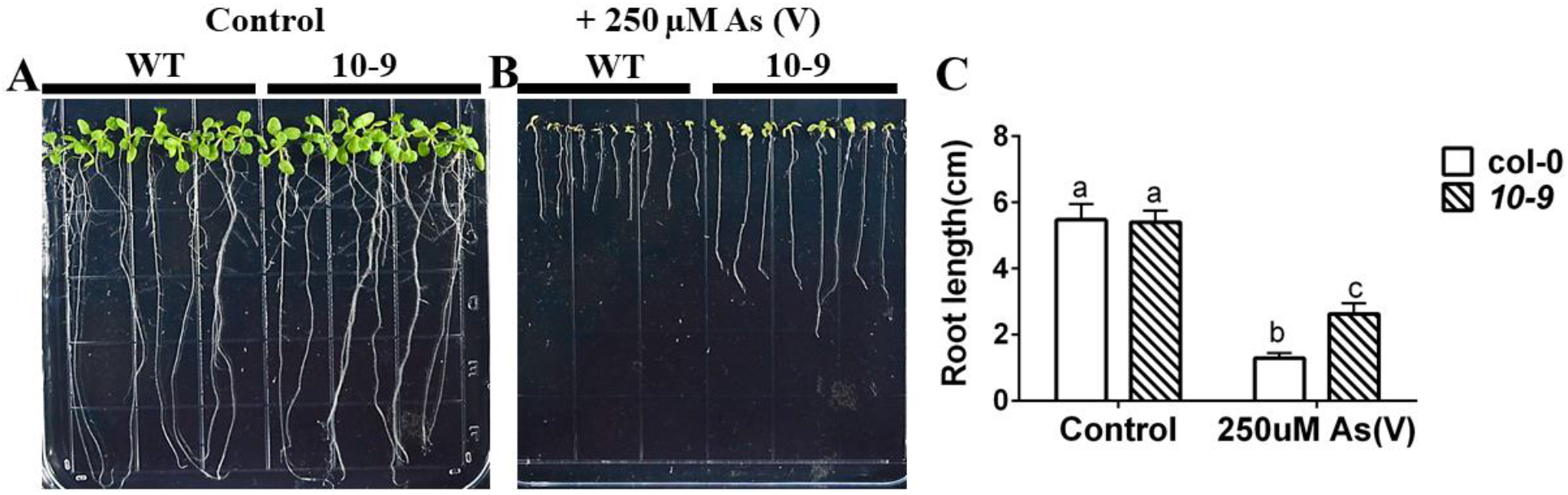
amiRNA *10-9* targets phosphate transporters and shows arsenate resistance. (A), (B) Root growth of amiRNA *10-9* is more tolerant than WT to arsenate treatment: WT and amiRNA *10-9* lines were grown vertically on 1/2 MS with 0.5% sucrose plates without (control) or with 250 μM sodium arsenate for 10 days; (C) Primary root lengths of WT and amiRNA *10-9* were significantly longer than WT on arsenite plates and no different on control plates. (n=15 seedlings, means ± s.e.m.). Letters at the top of columns are grouped based on two-way ANOVA and Tukey’s multiple comparisons test, P < 0.05).

The amiRNA *10-9* line was isolated and sequenced and is predicted to target three high-affinity phosphate transporters *(PHT1;1* AT5G43350, *PHT1;2* AT5G43370 and *PHT1;3* AT5G43360; amiRNA sequence listed in Table 1). In previous research, the PHT transporters AtPHT1;1, AtPHT1;4, AtPHT1;5; AtPHT1;7; 1;8; 1;9 were shown to function in both inorganic phosphate and arsenate (As(V)) transport in plants, as the oxyanion chemical structure of As(V) is structurally analogous to that of phosphate (Willsky & Malamy, 1980; Xu, Ma, & Nussinov, 2012). Interestingly however, the amiRNA line 10-9 also showed an enhanced sensitivity to arsenite (Figure 7A-C), which is not known to be a substrate of PHT phosphate transporters.

To determine whether the amiRNA *10-9* line phenotype is linked to the identified amiRNA, we amplified the original amiRNA sequence of amiRNA *10-9* and re-cloned and retransformed the amiRNA into Col-0 wild-type plants. The retransformed lines *Re-10-9-15* showed a decreased sensitivity to As(III) compared to WT controls in 5 of 10 independent transformants (Figure 7D-F). Note that this frequency of phenotypes upon amiRNA retransformation is not low and may be ascribed to the fact that amiRNAs partially inhibit transcription or translation at intermediate levels and phenotypes depend on individual transformation events (Schwab et al., 2006; Hauser et al., 2019). Thus, using amiRNA’s to target gene families, we have obtained evidence suggesting an unexpected role for PHT transporters in arsenite tolerance.

## Discussion

Forward genetic screens have enabled the identification of new genes and characterization of genetic pathways in *Arabidopsis* and other model organisms. While forward genetic screens continue to be powerful tools for unraveling biological processes, the presence of large gene families and many gene duplications with functional overlap in plant genomes, often buffers the effects of loss of function mutations leading to no observable phenotype in single-gene mutants (Lloyd & Meinke, 2012). Furthermore, higher-order loss of function mutations frequently lead to lethality, which also prevents the characterization of gene functions(Hauser et al., 2019). These pitfalls have hindered researchers’ ability to functionally characterize genes that belong to gene families (Lloyd & Meinke, 2012), which account for approximately 22,000 genes in *Arabidopsis*(Hauser et al., 2013).

To address these limitations, a library of artificial microRNAs (amiRNAs) was developed that targets diverse gene combinations mainly within sub-clades of multigene families to overcome genetic redundancy. This library has successfully enabled forward genetic screens to identify new genes involved in CO2 and ABA signaling as well as novel auxin transporters from very large gene families (Zhang et al., 2018; Hauser et al., 2019;Takahashi et al., 2020). Furthermore, because amiRNAs often cause transcriptional knockdown or translational inhibition rather than full loss of function, the amiRNA library has also enabled the identification of functionally redundant genes, for which double loss of function mutation causes lethality (Hauser et al., 2019). Together, these results demonstrate the power and utility of forward genetic screens using amiRNAs.

Arsenic exposure has been shown to induce a robust transcriptional response in *Arabidopsis*(Abercrombie et al., 2008; Jobe et al., 2012; Shukla et al., 2018). However, relatively few transcriptional regulators of arsenic-induced gene expression have been identified (Castrillo et al., 2013; Wang et al., 2014; Wang et al., 2017), likely due to extensive genetic redundancy in transcription factor families. Similarly, we hypothesized that additional transporters that affect arsenite sensitivity might exist, given the large family sizes and substrate overlap in plant transporters (Hauser et al., 2013). Thus, the genetic screen performed in the present study used amiRNA’s targeting DNA- and RNA-binding proteins and transporters and channels in *Arabidopsis*(Hauser et al., 2013). Notably, eight additional amiRNA libraries are available targeting other classes of proteins, including a library of over 4,000 amiRNAs targeting gene combinations within clades of gene families of presently unpredictable (“unknown”) function (Hauser et al., 2013;Hauser et al., 2019), suggesting that future screens using this platform are likely to uncover additional new genes that affect arsenic sensitivity.

In the present study, we identified the amiRNA *4-138* line as an As(III) insensitive line in root growth assays using a forward genetic screen. This amiRNA line targets the CBF1,2,3 and ERF34 and ERF35 transcription factors. We found that the As(III) insensitive phenotype is due to the tandem repeat genes *CBF1,2,3*. Note that *CBF4* may also contribute to this phenotype, but a *cbf* quadruple mutant may be lethal and was not investigated here. The *CBF1,2 and 3* genes are located in tandem on chromosome 4, and therefore CRSPR-mediated deletion of these genes, shows that they are not essential for seedling survival (Figure 1D).

By analyzing DAP-seq data sets for candidate gene promoters to which CBF1,2 & 3 bind (O’Malley et al., 2016), we identified 2214 genes potentially targeted by CBF1, CBF2, CBF3 and CBF4 (Supplemental Figure 6A and B). DAP-seq data suggest CBFs bind to many PHT transporter promoter regions. Further analyses showed that *PHT* transcript levels are decreased in *cbf1/2/3* triple mutants under non-arsenic stress control conditions.

The present findings suggest that CBF1,2,3 are positive regulators of PHT transporters. Additional *in-vivo* studies would allow further characterization of the underlying mechanisms.

Surprisingly, follow-up experiments with the arsenite tolerant amiRNA *4-138* line found this line to be cadmium sensitive. The contrasting root elongation phenotypes in response to arsenic (tolerant) and cadmium (sensitive) are unexpected as both cadmium and arsenite detoxification require thiolate peptides, in particular phytochelatins (Schmoger et al., 2000; Gong, Lee, & Schroeder, 2003; Aborode et al., 2016). As discussed above, the arsenite tolerant phenotype is due to the disruption of *CBF* genes. In contrast, characterization of the *erf34/35* double mutant suggests that the ERF transcription factors are the main contributors to the cadmium sensitivity phenotype of amiRNA *4-138.*

The nitrate transporter *NRT1.8* mRNA is one of the most strongly cadmium-induced transcripts based on microarray studies (Weber et al., 2006; Li et al., 2010). Furthermore, ERF34 and ERF35 transcription factors were found to bind the promoter of the NRT1.8 transporter in in-vitro binding assays (Table 3) (Weber et al., 2006; O’Malley et al., 2016). NRT1.8 is a plasma membrane transporter expressed predominantly in xylem parenchyma cells within the vasculature (Li et al., 2010). *NRT1.8* transcripts were strongly upregulated by cadmium stress in roots in the present study and a previous study (Li et al., 2010). Thus, the cadmium sensitive phenotype observed in *erf34/35* may be partly due to the misregulation of *NRT1.8.* The *nrt1.8* knock out mutant accumulates cadmium in shoots and has a similar root sensitivity to cadmium (Li et al., 2010) as the *erf34/35* double mutant (Figure 5). However, additional experiments are needed to confirm this hypothesis, as additional CBF/ERF targets are also likely involved.

Arsenate (As(V) can be transported by inorganic phosphate (Pi) transporters due to its chemical similarity to phosphate. Plasma membrane-localized Pi transporters are known to be responsible for arsenate uptake. In *Arabidopsis thaliana,* the As(V) transport capabilities of AtPHT1; 1, AtPHT1; 4, AtPHT1; 5, AtPHT1; 7, AtPHT1; 8, AtPHT1; 9 have been well characterized (Shin et al., 2004; Catarecha et al., 2007; Remy et al., 2012;LeBlanc et al., 2013; Fontenot et al., 2015). Decreased expression of these *PHT1* genes results in tolerance to As(V) (Shin et al., 2004; Catarecha et al., 2007; Nagarajan et al.,2011; LeBlanc et al., 2013; Luan et al., 2018). However, the involvement of PHT transporters in As(III) responses has not been previously established. In our study, amiRNA *10-9* seedlings showed reduced sensitivity to As(III) and As(V) in root elongation assays (Figures 1 & 7). AmiRNA *10-9* targets AtPHT1; 1, AtPHT1; 2, AtPHT1; 3, which share a high level of similarity, have overlapping expression patterns, and are reported to all contribute to Pi uptake (Ayadi et al., 2015). We have further found that both As(V) and As(III) downregulate the transcript levels of *AtPHT1; 1, AtPHT1; 2,* and *AtPHT1; 3*(Figures 3 & 4). Furthermore, PHT1 was shown to transport Pi and As(V) (Shin et al.,2004; Catarecha et al., 2007; Nussaume et al., 2011; Remy et al., 2012; LeBlanc et al.,2013; Fontenot et al., 2015). WRKY transcription factors, namely WRKY6, WRKY42, and WRKY45, were shown to interact with the PHT1;1 promoter and mediate *PHT1;1* expression (Castrillo et al., 2013; Wang et al., 2014; Su et al., 2015). WRKY6 and WRKY 42 were also shown to affect As(V) uptake into plants (Castrillo et al., 2013; Su et al.,2015), but As(III) sensitivity or transport were not analyzed in these mutants. Thus, the mechanism of As(III) resistance in the amiRNA *10-9* line is an unexpected interesting finding that requires further characterization.

In the present study, we pursued screening of amiRNA seed libraries that target diverse combinations of related DNA binding proteins and transporter proteins. Several putative amiRNA target genes, in addition to those characterized here were isolated. For example, we identified transcriptional regulators such as ethylene-response transcription factors, basic helix-loop-helix (bHLH) transcription factors, mitochondrial transcription termination factors, and ABCG transporters that warrant further investigation (Table 1).

In summary, the transcription factors that induce and repress gene expression in response to arsenic and cadmium remain to a large degree unknown. In the present study, a forward genetics screen of amiRNA lines that target combinatorial close homologs of DNA and RNA-binding proteins found that the CBF1, 2, and 3 transcription factors mediate negative regulation of arsenite and arsenate sensitivity and that the ERF34 and ERF35 transcription factors function in mediating cadmium resistance. Further studies of these transcription factors could identify the cadmium- and arsenic-induced transcriptional control network.

## Acknowledgments

This research was funded by the National Institute of Environmental Health Sciences of the National Institutes of Health under Award Number P42ES010337 (JIS) and in part by the National Natural Science Foundation of China (31770283) (SX).

## Figure Legends

**Supplemental Figure 1. amiRNA *4-85* targets ARR transcription factors, has long roots when grown on arsenite, and the phenotype was confirmed in amiRNA retransformation lines.**

(A), (B) amiRNA line *4-85* was more tolerant than WT under arsenite treatment: WT and *4-85* amiRNA lines were grown vertically on 1/2 MS with 0.5% sucrose plates without (control) or with 10 μM arsenite for 10 days; (C) Primary root lengths of amiRNA *4-85* compared to WT on control and arsenite-containing plates and on control plates. (n=15 seedlings, means ± s.e.m.). (D), (E) amiRNA retransformation line *4-85-10* are more tolerant than WT on germination assays: WT and amiRNA retransformation line *Re-4-85-10* were germinated directly on 1/2 MS with 0.5% sucrose plates without (control) or with 10 μM arsenite for about 14 days; (F) Primary root length of amiRNA *Re-4-85-10* compared to WT on control and arsenite-containing plates. (n=11 seedlings, means ± s.e.m.). Letters at the top of columns are grouped based on two-way ANOVA and Tukey’s multiple comparisons test, P < 0.05).

**Supplemental Figure 2. *erf34* and *erf35* single mutant shows weak or no phenotype under cadmium and arsenite treatment.**

(A) *erf34* and *erf35* has similar root lengths on control plates. (B) *erf35* has shorter roots compared to WT when Cd was added to the medium, but *erf34* shows no different when compared to WT; (C) Both *erf34* and *erf35* single mutants have no phenotype on arsenite containing plates: WT *erf34* and *erf35* single mutants were grown vertically on minimal plates with 1% sucrose plates without (control) or with 30 μM cadmium or with 10 μM sodium arsenite for 7 days; (D) (n=15 seedlings, means ± s.e.m.). Letters at the top of columns are grouped based on two-way ANOVA and Tukey’s multiple comparisons test, P < 0.05).

**Supplemental Figure 3. CRISPR/Cas9 gene editing technology to generate CRISPR *cbf1/2/3* knockout mutants**

(A) Cutting sites and primers for CRISPR *cbf1/2/3;* (B) Genotyping results for isolating homozygous for CRISPR *cbf1/2/3.*

**Supplemental Figure 4. Genotyping results for isolating homozygous for *erf34xerf35* knockout mutants**

(A) Genotyping results for *erf34* gene; (B) Genotyping results for *erf35* gene.

**Supplemental Figure 5. Venn diagram of genes targeted by ERF34, ERF35 and Genes significantly up regulated and downregulated after 2 hours of exposure to 50 μM Cd^2+^ in *A. thaliana* roots.**

**Supplemental Figure 6. Venn diagram and enriched ontology clusters of genes targeted by CBF1, 2, 3 and 4.**

(A) Venn diagrame; (B) Enriched ontology clusters of genes targed by CBF1, 2, 3, 4 in the same time.

## Table Legends

**Supplemental Table 1. Primers used for amiRNA sequencing and cloning**

**Supplemental Table 2. Salk lines and primers for genotyping**

**Supplemental Table 3. Primers used for RTq-PCR**

## References

Abbas, G., Murtaza, B., Bibi, I., Shahid, M., Niazi, N. K., Khan, M. I.,… Natasha. (2018). Arsenic Uptake, Toxicity, Detoxification, and Speciation in Plants: Physiological, Biochemical, and Molecular Aspects. Int J Environ Res Public Health, 15(1). doi:10.3390/ijerph15010059

Abedi, T., & Mojiri, A. (2020). Arsenic Uptake and Accumulation Mechanisms in Rice Species. Plants (Basel), 9(2). doi:10.3390/plants9020129

Abercrombie, J. M., Halfhill, M. D., Ranjan, P., Rao, M. R., Saxton, A. M., Yuan, J. S., & Stewart, C. N., Jr. (2008). Transcriptional responses of Arabidopsis thaliana plants to As (V) stress. BMC Plant Biol, 8, 87. doi:10.1186/1471-2229-8-87

Aborode, F. A., Raab, A., Voigt, M., Costa, L. M., Krupp, E. M., & Feldmann, J. (2016). The importance of glutathione and phytochelatins on the selenite and arsenate detoxification in Arabidopsis thaliana. J Environ Sci (China), 49, 150–161. doi:10.1016/j.jes.2016.08.009

Arabidopsis Genome, I. (2000). Analysis of the genome sequence of the flowering plant Arabidopsis thaliana. Nature, 408(6814), 796–815. doi:10.1038/35048692

Ayadi, A., David, P., Arrighi, J. F., Chiarenza, S., Thibaud, M. C., Nussaume, L., & Marin, E. (2015). Reducing the genetic redundancy of Arabidopsis PHOSPHATE TRANSPORTER1 transporters to study phosphate uptake and signaling. Plant Physiol, 167(4), 1511–1526. doi:10.1104/pp.114.252338

Bienert, G. P., Thorsen, M., Schussler, M. D., Nilsson, H. R., Wagner, A., Tamas, M. J., & Jahn, T. P. (2008). A subgroup of plant aquaporins facilitate the bi-directional diffusion of As(OH)3 and Sb(OH)3 across membranes. BMC Biol, 6, 26. doi:10.1186/1741-7007-6-26

Castrillo, G., Sanchez-Bermejo, E., de Lorenzo, L., Crevillen, P., Fraile-Escanciano, A., Tc, M.,… Leyva, A. (2013). WRKY6 transcription factor restricts arsenate uptake and transposon activation in Arabidopsis. Plant Cell, 25(8), 2944–2957. doi:10.1105/tpc.113.114009

Catarecha, P., Segura, M. D., Franco-Zorrilla, J. M., Garcia-Ponce, B., Lanza, M., Solano, R., & Leyva, A. (2007). A mutant of the Arabidopsis phosphate transporter PHT1;1 displays enhanced arsenic accumulation. Plant Cell, 19(3), 1123–1133. doi:10.1105/tpc.106.041871

Chakrabarty, D., Trivedi, P. K., Misra, P., Tiwari, M., Shri, M., Shukla, D., & Tuli, R. (2009). Comparative transcriptome analysis of arsenate and arsenite stresses in rice seedlings. Chemosphere, 74(5), 688–702. doi:10.1016/j.chemosphere.2008.09.082

Chao, D. Y., Chen, Y., Chen, J., Shi, S., Chen, Z., Wang, C., & Salt, D. E. (2014). Genome-wide association mapping identifies a new arsenate reductase enzyme critical for limiting arsenic accumulation in plants. PLoS Biol, 12(12), e1002009. doi:10.1371/journal.pbio.1002009

Clemens, S. (2019). Safer food through plant science: reducing toxic element accumulation in crops. J Exp Bot, 70(20), 5537–5557. doi:10.1093/jxb/erz366

Clemens, S., & Ma, J. F. (2016). Toxic Heavy Metal and Metalloid Accumulation in Crop Plants and Foods. Annu Rev Plant Biol, 67, 489–512. doi:10.1146/annurev-arplant-043015-112301

Clough, S. J., & Bent, A. F. (1998). Floral dip: a simplified method for Agrobacterium-mediated transformation of Arabidopsis thaliana. Plant J, 16(6), 735–743. doi:10.1046/j.1365-313x.1998.00343.x

Cooper, A. M., Felix, D., Alcantara, F., Zaslavsky, I., Work, A., Watson, P. L., & Schroeder, J. I. (2020). Monitoring and mitigation of toxic heavy metals and arsenic accumulation in food crops: A case study of an urban community garden. Plant Direct, 4(1), e00198. doi:10.1002/pld3.198

Cusack, S. A., Wang, P., Moore, B. M., Meng, F., Conner, J. K., Krysan, P. J., & Shiu, S.-H. (2020). Genome-wide predictions of genetic redundancy in Arabidopsis thaliana. bioRxiv.

Duan, G. L., Hu, Y., Schneider, S., McDermott, J., Chen, J., Sauer, N., & Zhu, Y. G. (2016a). Inositol transporters AtINT2 and AtINT4 regulate arsenic accumulation in Arabidopsis seeds. Nature Plants, 2(1). doi:Artn 15202 10.1038/Nplants.2015.202

Duan, G. L., Hu, Y., Schneider, S., McDermott, J., Chen, J., Sauer, N., & Zhu, Y. G. (2016b). Inositol transporters AtINT2 and AtINT4 regulate arsenic accumulation in Arabidopsis seeds. Nat Plants, 2(1), 15202. doi:10.1038/nplants.2015.202

Fontenot, E. B., Ditusa, S. F., Kato, N., Olivier, D. M., Dale, R., Lin, W. Y., & Smith, A. P. (2015). Increased phosphate transport of Arabidopsis thaliana Pht1;1 by site-directed mutagenesis of tyrosine 312 may be attributed to the disruption of homomeric interactions. Plant Cell Environ, 38(10), 2012–2022. doi:10.1111/pce.12522

Gao, X., Chen, J., Dai, X., Zhang, D., & Zhao, Y. (2016). An Effective Strategy for Reliably Isolating Heritable and Cas9-Free Arabidopsis Mutants Generated by CRISPR/Cas9-Mediated Genome Editing. Plant Physiol, 171(3), 1794–1800. doi:10.1104/pp.16.00663

Gao, Y., & Zhao, Y. (2014). Self-processing of ribozyme-flanked RNAs into guide RNAs in vitro and in vivo for CRISPR-mediated genome editing. J Integr Plant Biol, 56(4), 343–349. doi:10.1111/jipb.12152

Gao, Y., Zhang, Y., Zhang, D., Dai, X., Estelle, M., & Zhao, Y. (2015). Auxin binding protein 1 (ABP1) is not required for either auxin signaling or Arabidopsis development. Proc Natl Acad Sci USA, 112(7), 2275–2280. doi:10.1073/pnas.1500365112

Garbinski, L. D., Rosen, B. P., & Chen, J. (2019). Pathways of arsenic uptake and efflux. Environ Int, 126, 585–597. doi:10.1016/j.envint.2019.02.058

Gasic, K., & Korban, S. S. (2007). Transgenic Indian mustard (Brassica juncea) plants expressing an Arabidopsis phytochelatin synthase (AtPCS1) exhibit enhanced As and Cd tolerance. Plant Molecular Biology, 64(4), 361–369. doi:10.1007/s11103-007-9158-7

Gilbert-Diamond, D., Cottingham, K. L., Gruber, J. F., Punshon, T., Sayarath, V., Gandolfi, A. J., & Karagas, M. R. (2011). Rice consumption contributes to arsenic exposure in US women. Proc Natl Acad Sci USA, 108(51), 20656–20660. doi:10.1073/pnas.1109127108

Gong, J. M., Lee, D. A., & Schroeder, J. I. (2003). Long-distance root-to-shoot transport of phytochelatins and cadmium in Arabidopsis. Proc Natl Acad Sci U S A, 100(17), 10118–10123.

Guo, B., Jin, Y., Wussler, C., Blancaflor, E. B., Motes, C. M., & Versaw, W. K. (2008). Functional analysis of the Arabidopsis PHT4 family of intracellular phosphate transporters. New Phytol, 177(4), 889–898. doi:10.1111/j.1469-8137.2007.02331.x

Guo, J., Dai, X., Xu, W., & Ma, M. (2008). Overexpressing GSH1 and AsPCS1 simultaneously increases the tolerance and accumulation of cadmium and arsenic in Arabidopsis thaliana. Chemosphere, 72(7), 1020–1026. doi:10.1016/j.chemosphere.2008.04.018

Hauser, F., Chen, W., Deinlein, U., Chang, K., Ossowski, S., Fitz, J., & Schroeder, J. I. (2013). A genomic-scale artificial microRNA library as a tool to investigate the functionally redundant gene space in Arabidopsis. Plant Cell, 25(8), 2848–2863. doi:10.1105/tpc.113.112805

Hauser, F., Ceciliato, P. H. O., Lin, Y. C., Guo, D., Gregerson, J. D., Abbasi, N., & Schroeder, J. I. (2019). A seed resource for screening functionally redundant genes and isolation of new mutants impaired in CO2 and ABA responses. J Exp Bot, 70(2), 641–651. doi:10.1093/jxb/ery363

Hayashi, S., Kuramata, M., Abe, T., Takagi, H., Ozawa, K., & Ishikawa, S. (2017). Phytochelatin synthase OsPCS1 plays a crucial role in reducing arsenic levels in rice grains. Plant Journal, 91(5), 840–848. doi:10.1111/tpj.13612

Herschbach, C., Rizzini, L., Mult, S., Hartmann, T., Busch, F., Peuke, A. D., & Ensminger, I. (2010). Over-expression of bacterial gamma-glutamylcysteine synthetase (GSH1) in plastids affects photosynthesis, growth and sulphur metabolism in poplar (Populus tremula x Populus alba) dependent on the resulting gamma-glutamylcysteine and glutathione levels. Plant Cell Environ, 33(7), 1138–1151. doi:10.1111/j.1365-3040.2010.02135.x

Huang, Y., Chen, H., Reinfelder, J. R., Liang, X., Sun, C., Liu, C., & Yi, J. (2019). A transcriptomic (RNA-seq) analysis of genes responsive to both cadmium and arsenic stress in rice root. Sci Total Environ, 666, 445–460. doi:10.1016/j.scitotenv.2019.02.281

Isayenkov, S. V., & Maathuis, F. J. (2008). The Arabidopsis thaliana aquaglyceroporin AtNIP7;1 is a pathway for arsenite uptake. FEBS Lett, 582(11), 1625–1628. doi:10.1016/j.febslet.2008.04.022

Ji, R., Zhou, L., Liu, J., Wang, Y., Yang, L., Zheng, Q., & Lan, W. (2017). Calcium-dependent protein kinase CPK31 interacts with arsenic transporter AtNIP1;1 and regulates arsenite uptake in Arabidopsis thaliana. PLoS One, 12(3), e0173681. doi:10.1371/journal.pone.0173681

Jobe, T. O., Sung, D. Y., Akmakjian, G., Pham, A., Komives, E. A., Mendoza-Cozatl, D. G., & Schroeder, J. I. (2012). Feedback inhibition by thiols outranks glutathione depletion: a luciferase-based screen reveals glutathione-deficient gamma-ECS and glutathione synthetase mutants impaired in cadmium-induced sulfate assimilation. Plant J, 70(5), 783–795. doi:10.1111/j.1365-313X.2012.04924.x

Kamiya, T., & Fujiwara, T. (2009). Arabidopsis NIP1;1 transports antimonite and determines antimonite sensitivity. Plant Cell Physiol, 50(11), 1977–1981. doi:10.1093/pcp/pcp130

Kamiya, T., Tanaka, M., Mitani, N., Ma, J. F., Maeshima, M., & Fujiwara, T. (2009). NIP1;1, an aquaporin homolog, determines the arsenite sensitivity of Arabidopsis thaliana. J Biol Chem, 284(4), 2114–2120. doi:10.1074/jbc.M806881200

Kang, H. W., Cho, Y. G., Yoon, U. H., & Eun, M. Y. (1998). A Rapid DNA Extraction Method for RFLP and PCR Analysis from a Single Dry Seed. Plant Molecular Biology Reporter, 16(1), 90–90. doi:10.1023/A:1007418606098

Kumari, P., Rastogi, A., Shukla, A., Srivastava, S., & Yadav, S. (2018). Prospects of genetic engineering utilizing potential genes for regulating arsenic accumulation in plants. Chemosphere, 211, 397–406. doi:10.1016/j.chemosphere.2018.07.152

Lapis-Gaza, H. R., Jost, R., & Finnegan, P. M. (2014). Arabidopsis PHOSPHATE TRANSPORTER1 genes PHT1;8 and PHT1;9 are involved in root-to-shoot translocation of orthophosphate. BMC Plant Biol, 14, 334. doi:10.1186/s12870-014-0334-z

LeBlanc, M. S., McKinney, E. C., Meagher, R. B., & Smith, A. P. (2013). Hijacking membrane transporters for arsenic phytoextraction. J Biotechnol, 163(1), 1–9. doi:10.1016/j.jbiotec.2012.10.013

Lee, D. A., Chen, A., & Schroeder, J. I. (2003). ars1, an Arabidopsis mutant exhibiting increased tolerance to arsenate and increased phosphate uptake. Plant J, 35(5), 637–646. doi:10.1046/j.1365-313x.2003.01835.x

Li, J. Y., Fu, Y. L., Pike, S. M., Bao, J., Tian, W., Zhang, Y., & Gong, J. M. (2010). The Arabidopsis nitrate transporter NRT1.8 functions in nitrate removal from the xylem sap and mediates cadmium tolerance. Plant Cell, 22(5), 1633–1646. doi:10.1105/tpc.110.075242

Li, Y., Dhankher, O. P., Carreira, L., Lee, D., Chen, A., Schroeder, J. I., & Meagher, R. B. (2004). Overexpression of phytochelatin synthase in Arabidopsis leads to enhanced arsenic tolerance and cadmium hypersensitivity. Plant Cell Physiol, 45(12), 1787–1797. doi:10.1093/pcp/pch202

Lindsay, E. R., & Maathuis, F. J. (2016). Arabidopsis thaliana NIP7;1 is involved in tissue arsenic distribution and tolerance in response to arsenate. FEBS Lett, 590(6), 779–786. doi:10.1002/1873-3468.12103

Lindsay, E. R., & Maathuis, F. J. M. (2017). New Molecular Mechanisms to Reduce Arsenic in Crops. Trends in Plant Science, 22(12), 1016–1026. doi:10.1016/j.tplants.2017.09.015

Liu, Q. P., Hu, H. C., Zhu, L. Y., Li, R. C., Feng, Y., Zhang, L. Q., & Zhang, H. M. (2015). Involvement of miR528 in the Regulation of Arsenite Tolerance in Rice (Oryza sativa L.). Journal of Agricultural and Food Chemistry, 63(40), 8849–8861. doi:10.1021/acs.jafc.5b04191

Liu, X., Zhang, S., Shan, X., & Zhu, Y. G. (2005). Toxicity of arsenate and arsenite on germination, seedling growth and amylolytic activity of wheat. Chemosphere, 61(2), 293–301. doi:10.1016/j.chemosphere.2005.01.088

Lloyd, J., & Meinke, D. (2012). A comprehensive dataset of genes with a loss-of-function mutant phenotype in Arabidopsis. Plant Physiol, 158(3), 1115–1129. doi:10.1104/pp.111.192393

Luan, M., Liu, J., Liu, Y., Han, X., Sun, G., Lan, W., & Luan, S. (2018). Vacuolar Phosphate Transporter 1 (VPT1) Affects Arsenate Tolerance by Regulating Phosphate Homeostasis in Arabidopsis. Plant Cell Physiol, 59(7), 1345–1352. doi:10.1093/pcp/pcy025

Mendoza-Cozatl, D. G., Jobe, T. O., Hauser, F., & Schroeder, J. I. (2011). Long-distance transport, vacuolar sequestration, tolerance, and transcriptional responses induced by cadmium and arsenic. Curr Opin Plant Biol, 14(5), 554–562. doi:10.1016/j.pbi.2011.07.004

Mendoza-Cozatl, D. G., Xie, Q., Akmakjian, G. Z., Jobe, T. O., Patel, A., Stacey, M. G., & Schroeder, J. I. (2014). OPT3 is a component of the iron-signaling network between leaves and roots and misregulation of OPT3 leads to an over-accumulation of cadmium in seeds. Mol Plant, 7(9), 1455–1469. doi:10.1093/mp/ssu067

Mosa, K. A., Kumar, K., Chhikara, S., McDermott, J., Liu, Z., Musante, C., & Dhankher, O. P. (2012). Members of rice plasma membrane intrinsic proteins subfamily are involved in arsenite permeability and tolerance in plants. Transgenic Res, 21(6), 1265–1277. doi:10.1007/s11248-012-9600-8

Nagarajan, V. K., Jain, A., Poling, M. D., Lewis, A. J., Raghothama, K. G., & Smith, A. P. (2011). Arabidopsis Pht1;5 mobilizes phosphate between source and sink organs and influences the interaction between phosphate homeostasis and ethylene signaling. Plant Physiol, 156(3), 1149–1163. doi:10.1104/pp.111.174805

Nussaume, L., Kanno, S., Javot, H., Marin, E., Pochon, N., Ayadi, A., & Thibaud, M. C. (2011). Phosphate Import in Plants: Focus on the PHT1 Transporters. Front Plant Sci, 2, 83. doi:10.3389/fpls.2011.00083

O’Malley, R. C., Huang, S. C., Song, L., Lewsey, M. G., Bartlett, A., Nery, J. R., & Ecker, J. R. (2016). Cistrome and Epicistrome Features Shape the Regulatory DNA Landscape. Cell, 166(6), 1598. doi:10.1016/j.cell.2016.08.063

Ossowski, S., Fitz, J., Schwab, R., Riester, M., & Weigel, D. personal communication.

Palma-Lara, I., Martinez-Castillo, M., Quintana-Perez, J. C., Arellano-Mendoza, M. G., Tamay-Cach, F., Valenzuela-Limon, O. L., & Hernandez-Zavala, A. (2020). Arsenic exposure: A public health problem leading to several cancers. Regul Toxicol Pharmacol, 110, 104539. doi:10.1016/j.yrtph.2019.104539

Pan, W., Wu, C., Xue, S., & Hartley, W. (2014). Arsenic dynamics in the rhizosphere and its sequestration on rice roots as affected by root oxidation. J Environ Sci (China), 26(4), 892–899. doi:10.1016/S1001-0742(13)60483-0

Picault, N., Cazale, A. C., Beyly, A., Cuine, S., Carrier, P., Luu, D. T., & Peltier, G. (2006). Chloroplast targeting of phytochelatin synthase in Arabidopsis: effects on heavy metal tolerance and accumulation. Biochimie, 88(11), 1743–1750. doi:10.1016/j.biochi.2006.04.016

Qian, L., Qi, S., Cao, F., Zhang, J., Zhao, F., Li, C., & Wang, C. (2018). Toxic effects of boscalid on the growth, photosynthesis, antioxidant system and metabolism of Chlorella vulgaris. Environ Pollut, 242(Pt A), 171–181. doi:10.1016/j.envpol.2018.06.055

Rai, P. K., Lee, S. S., Zhang, M., Tsang, Y. F., & Kim, K. H. (2019). Heavy metals in food crops: Health risks, fate, mechanisms, and management. Environ Int, 125, 365–385. doi:10.1016/j.envint.2019.01.067

Remy, E., Cabrito, T. R., Batista, R. A., Teixeira, M. C., Sa-Correia, I., & Duque, P. (2012). The Pht1;9 and Pht1;8 transporters mediate inorganic phosphate acquisition by the Arabidopsis thaliana root during phosphorus starvation. New Phytol, 195(2), 356–371. doi:10.1111/j.1469-8137.2012.04167.x

Schmoger, M. E., Oven, M., & Grill, E. (2000). Detoxification of arsenic by phytochelatins in plants. Plant Physiol, 122(3), 793–801. doi:10.1104/pp.122.3.793

Schwab, R., Ossowski, S., Riester, M., Warthmann, N., & Weigel, D. (2006). Highly specific gene silencing by artificial microRNAs in Arabidopsis. Plant Cell, 18(5), 1121–1133. doi:10.1105/tpc.105.039834

Shi, S., Wang, T., Chen, Z., Tang, Z., Wu, Z., Salt, D. E., & Zhao, F. J. (2016). OsHAC1;1 and OsHAC1;2 Function as Arsenate Reductases and Regulate Arsenic Accumulation. Plant Physiol, 172(3), 1708–1719. doi:10.1104/pp.16.01332

Shin, H., Shin, H. S., Dewbre, G. R., & Harrison, M. J. (2004). Phosphate transport in Arabidopsis: Pht1;1 and Pht1;4 play a major role in phosphate acquisition from both low- and high-phosphate environments. Plant J, 39(4), 629–642. doi:10.1111/j.1365-313X.2004.02161.x

Shri, M., Singh, P. K., Kidwai, M., Gautam, N., Dubey, S., Verma, G., & Chakrabarty, D. (2019). Recent advances in arsenic metabolism in plants: current status, challenges and highlighted biotechnological intervention to reduce grain arsenic in rice. Metallomics, 11(3), 519–532. doi:10.1039/c8mt00320c

Shukla, T., Khare, R., Kumar, S., Lakhwani, D., Sharma, D., Asif, M. H., & Trivedi, P. K. (2018). Differential transcriptome modulation leads to variation in arsenic stress response in Arabidopsis thaliana accessions. J Hazard Mater, 351, 1–10. doi:10.1016/j.jhazmat.2018.02.031

Song, W. Y., Yamaki, T., Yamaji, N., Ko, D., Jung, K. H., Fujii-Kashino, M., & Ma, J. F. (2014). A rice ABC transporter, OsABCC1, reduces arsenic accumulation in the grain. Proceedings of the National Academy of Sciences of the United States of America, 111(44), 15699–15704. doi:10.1073/pnas.1414968111

Song, W. Y., Park, J., Mendoza-Cozatl, D. G., Suter-Grotemeyer, M., Shim, D., Hortensteiner, S., & Martinoia, E. (2010). Arsenic tolerance in Arabidopsis is mediated by two ABCC-type phytochelatin transporters. Proceedings of the National Academy of Sciences of the United States of America, 107(49), 21187–21192. doi:10.1073/pnas.1013964107

Srivastava, S., Srivastava, A. K., Sablok, G., Deshpande, T. U., & Suprasanna, P. (2015). Transcriptomics profiling of Indian mustard (Brassica juncea) under arsenate stress identifies key candidate genes and regulatory pathways. Front Plant Sci, 6, 646. doi:10.3389/fpls.2015.00646

Su, T., Xu, Q., Zhang, F. C., Chen, Y., Li, L. Q., Wu, W. H., & Chen, Y. F. (2015). WRKY42 modulates phosphate homeostasis through regulating phosphate translocation and acquisition in Arabidopsis. Plant Physiol, 167(4), 1579–1591. doi:10.1104/pp.114.253799

Takahashi, Y., Zhang, J., Hsu, P. K., Ceciliato, P. H. O., Zhang, L., Dubeaux, G., & Schroeder, J. I. (2020). MAP3Kinase-dependent SnRK2-kinase activation is required for abscisic acid signal transduction and rapid osmotic stress response. Nat Commun, 11(1), 12. doi:10.1038/s41467-019-13875-y

Tiwari, M., Sharma, D., Dwivedi, S., Singh, M., Tripathi, R. D., & Trivedi, P. K. (2014). Expression in Arabidopsis and cellular localization reveal involvement of rice NRAMP, OsNRAMP1, in arsenic transport and tolerance. Plant Cell Environ, 37(1), 140–152. doi:10.1111/pce.12138

Verma, P. K., Verma, S., Pande, V., Mallick, S., Tripathi, R. D., Dhankher, O. P., & Chakrabarty, D. (2017). Overexpression of Rice Glutaredoxin OsGrx_C7 and OsGrx_C2.1 Reduces Intracellular Arsenic Accumulation and Increases Tolerance in Arabidopsis thaliana (vol 7, 740, 2016). Frontiers in Plant Science, 8. doi:ARTN 1884 10.3389/fpls.2017.01884

Versaw, W. K., & Harrison, M. J. (2002). A chloroplast phosphate transporter, PHT2;1, influences allocation of phosphate within the plant and phosphate-starvation responses. Plant Cell, 14(8), 1751–1766. doi:10.1105/tpc.002220

Wang, F. Z., Chen, M. X., Yu, L. J., Xie, L. J., Yuan, L. B., Qi, H., & Chen, Q. F. (2017). OsARM1, an R2R3 MYB Transcription Factor, Is Involved in Regulation of the Response to Arsenic Stress in Rice. Front Plant Sci, 8, 1868. doi:10.3389/fpls.2017.01868

Wang, H., Xu, Q., Kong, Y. H., Chen, Y., Duan, J. Y., Wu, W. H., & Chen, Y. F. (2014). Arabidopsis WRKY45 transcription factor activates PHOSPHATE TRANSPORTER1;1 expression in response to phosphate starvation. Plant Physiol, 164(4), 2020–2029. doi:10.1104/pp.113.235077

Weber, M., Trampczynska, A., & Clemens, S. (2006). Comparative transcriptome analysis of toxic metal responses in Arabidopsis thaliana and the Cd(2+)-hypertolerant facultative metallophyte Arabidopsis halleri. Plant Cell Environ, 29(5), 950–963. doi:10.1111/j.1365-3040.2005.01479.x

Willsky, G. R., & Malamy, M. H. (1980). Effect of arsenate on inorganic phosphate transport in Escherichia coli. J Bacteriol, 144(1), 366–374. doi:10.1128/JB.144.1.366-374.1980

Wojas, S., Clemens, S., Sklodowska, A., & Maria Antosiewicz, D. (2010). Arsenic response of AtPCS1- and CePCS-expressing plants - effects of external As(V) concentration on As-accumulation pattern and NPT metabolism. J Plant Physiol, 167(3), 169–175. doi:10.1016/j.jplph.2009.07.017

Xu, J. M., Shi, S. L., Wang, L., Tang, Z., Lv, T. T., Zhu, X. L., & Wu, Z. C. (2017). OsHAC4 is critical for arsenate tolerance and regulates arsenic accumulation in rice. New Phytologist, 215(3), 1090–1101. doi:10.1111/nph.14572

Xu, W., Dai, W., Yan, H., Li, S., Shen, H., Chen, Y., & Ma, M. (2015). Arabidopsis NIP3;1 Plays an Important Role in Arsenic Uptake and Root-to-Shoot Translocation under Arsenite Stress Conditions. Mol Plant, 8(5), 722–733. doi:10.1016/j.molp.2015.01.005

Xu, Y., Ma, B., & Nussinov, R. (2012). Structural and functional consequences of phosphate-arsenate substitutions in selected nucleotides: DNA, RNA, and ATP. J Phys Chem B, 116(16), 4801–4811. doi:10.1021/jp300307u

Yang, H., Menz, J., Haussermann, I., Benz, M., Fujiwara, T., & Ludewig, U. (2015). High and Low Affinity Urea Root Uptake: Involvement of NIP5;1. Plant Cell Physiol, 56(8), 1588–1597. doi:10.1093/pcp/pcv067

Yoon, Y., Lee, W. M., & An, Y. J. (2015). Phytotoxicity of arsenic compounds on crop plant seedlings. Environ Sci Pollut Res Int, 22(14), 11047–11056. doi:10.1007/s11356-015-4317-x

Zhang, Y., Nasser, V., Pisanty, O., Omary, M., Wulff, N., Di Donato, M., & Shani, E. (2018). A transportome-scale amiRNA-based screen identifies redundant roles of Arabidopsis ABCB6 and ABCB20 in auxin transport. Nat Commun, 9(1), 4204. doi:10.1038/s41467-018-06410-y

Zvobgo, G., Sagonda, T., Lwalaba, J. L. W., Mapodzeke, J. M., Muhammad, N., Chen, G., & Zhang, G. (2018). Transcriptomic comparison of two barley genotypes differing in arsenic tolerance exposed to arsenate and phosphate treatments. Plant Physiol Biochem, 130, 589–603. doi:10.1016/j.plaphy.2018.08.006

