## Supplemental data 1 for "An amiRNA screen uncovers redundant CBF & ERF34/35 transcription factors that differentially regulate arsenite and cadmium responses"

#### Supplemental Table 1. Primers used for amiRNA sequencing and cloning

| PRIMER NAME | SEQUENCE |
|---|---|
| **PHA2804F** | AGAGAACACGGGGGACGAG |
| **PHA3479R** | AAACCGGCGGTAAGGATCTG |
| **RE-AMI-F** | GGGGACAAGTTTGTACAAAAAAGCAGGCTTAGAATTCCTGCAGCCCCAAACACACG |
| **RE-AMI-R** | GGGGACCACTTTGTACAAGAAAGCTGGGTCGGATCCCCCCATGGCGATGCCTTA |

#### Supplemental Table 2. Salk lines and primers for genotyping

| GENE | GENE ID | SALK LINE | PRIMER SEQUENCE |
|---|---|---|---|
| **ERF34** | AT2G44940 | SALK_020979C | LB1.3: ATTTTGCCGATTTCGGAAC |
| | | | LP: TGTTAAGAGGTGACGCACATG |
| | | | RP: ACTCACATCGTTGACCGAATC |
| **ERF35** | AT3G60490 | SALK_111486C | LB1.3: ATTTTGCCGATTTCGGAAC |
| | | | LP: TTATCGATAACCGGTTTGTGC |
| | | | RP: TAAAACAATCCAGACCCATGC |
| **CRISP RCBF1/ 2/3** | AT4G25490 | GT1 | AGACAGCGAGTGGAACATCG |
| | AT4G25470 | GT2 | TTGTTGCTTATGGGGAGACCA |
| | AT4G25490 | GT3 | AAGTCCCGAGCCAAATCCTG |

#### Supplemental Table 3. Primers used for RTq-PCR

| PRRMER | SEQUENCE |
|---|---|
| **PHT1;1-QF** | GTTCTTAATTTCTCCTGCCAAGCTGATTA |
| **PHT1;1-QR** | CATCATAACTTAAGGTCAACGAGCCAATA |
| **PHT1;2-QF** | GAACAACAACTAGGAGTGCTAAAGGCACT |
| **PHT1;2-QR** | CTAACTTCAGCTTCACCAGAGAGTTCTTC |
| **PHT1;3-QF** | GATCAACAGCTAGGAGTGCTAAAGGCACT |
| **PHT1;3-QR** | TATCAACCTCAGCCTCGCCGGAGAGTTCT |
| **PHT1;4-QF** | CTCTCTCAACATTTTCCCCTGAAAATAAG |
| **PHT1;4-QR** | AGAACAACTTGAGTTGCTAGAGACAAGGA |
| **PHT1;5-QF** | ACGCAGCCAAAACGCAGATGTACCATTTC |
| **PHT1;5-QR** | GGACTTTTCTACCGGAATTTGCCACAGTC |
| **PHT1;6-QF** | ACGTTATACATCATGGCAGGAATCAAT |
| **PHT1;6-QR** | AAGCTCCTCAAGTGATTTCCCATTAGT |
| **PHT1;7-QF** | TGGAGGATATCCATGCTCTGTCT |
| **PHT1;7-QR** | CGCGGCTTCTGGAAAATTAG |
| **PHT1;8-QF** | GCTCTTCCTGCTGCATTGACG |
| **PHT1;8-QR** | AGGAGGTGGATCCGTCGTGGCTT |
| **PHT1;9-QF** | CCGCCAGATACACAGCATTGGT |
| **PHT1;9-QR** | CGACCGTGGAGACTGAGGAAAC |
| **PHT4;1-QF** | TCTTCTGGGGTTACCTTCTTACACAGA |
| **PHT4;1-QR** | TGAGAATTGTAGCGATTGACCACCAA |
| **PHT4;2-QF** | GATGATGCCTGAGAGGATTAAGGTAGT |
| **PHT4;2-QR** | TAACAACTCTGTCGGCGTTACATAGA |
| **PHT4;3-QF** | CAGAATTTATAACGTCGGAGAGAGTCAAA |
| **PHT4;3-QR** | AAAATGATTTGCTCCATCCACGAGAAAG |
| **PHT4;4-QF** | TGTGAACATGAGCATTGCAATTCTT |
| **PHT4;4-QR** | CAACAGTTGCACTACTCCAGTTATATTCT |
| **PHT4;5-QF** | TCCAGTCTTCCTTCTTTTGGGGTTATG |
| **PHT4;5-QR** | CGAAAGACCATGTAAAGACACCAATCT |
| **PHT4;6-QF** | GATTGGTTTCGATAACGACATCAGGAA |
| **PHT4;6-QR** | ATTCCGGACCTCTAAGCTCAACTAA |
| **PHT2;1-QF** | CTCCTCTCTTCCTTAGCTGCAGCTG |
| **PHT2;1-QR** | GCGAATGACATGAAGCAAGCGGAGAG |
| **HP30-1-QF** | GCGTTGACCCTCAAGCCATA |
| **HP30-1-QR** | CTGCCACCACTGCAGATTCA |
| **HP30-2-QF** | TTAGCTGACCCAACTCTGCC |
| **HP30-2-QR** | TAGCTCAGGGTCCCTTTGGA |
| **CPK31-QF** | CGCTGGGAGTGCTTACTACATTGC |
| **CPK31-QR** | ACTTCCATGATTCGCTGTCAACGTC |

### Supplemental Figure 1

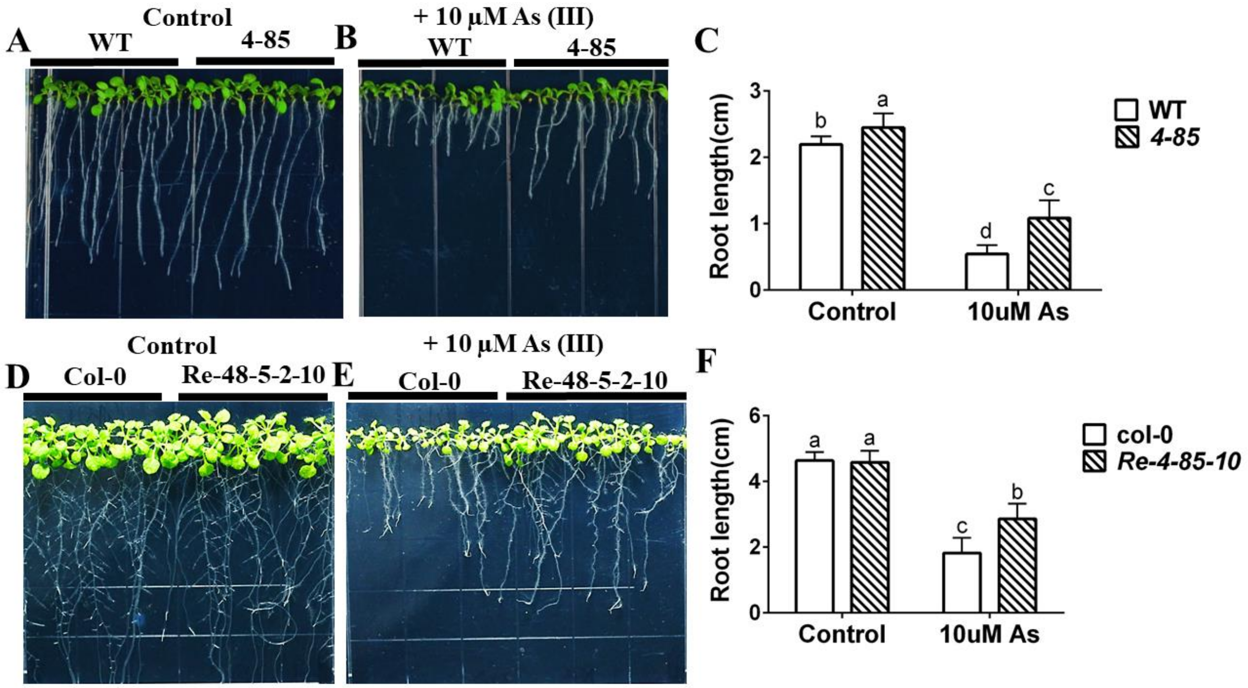

#### Supplemental Figure 2

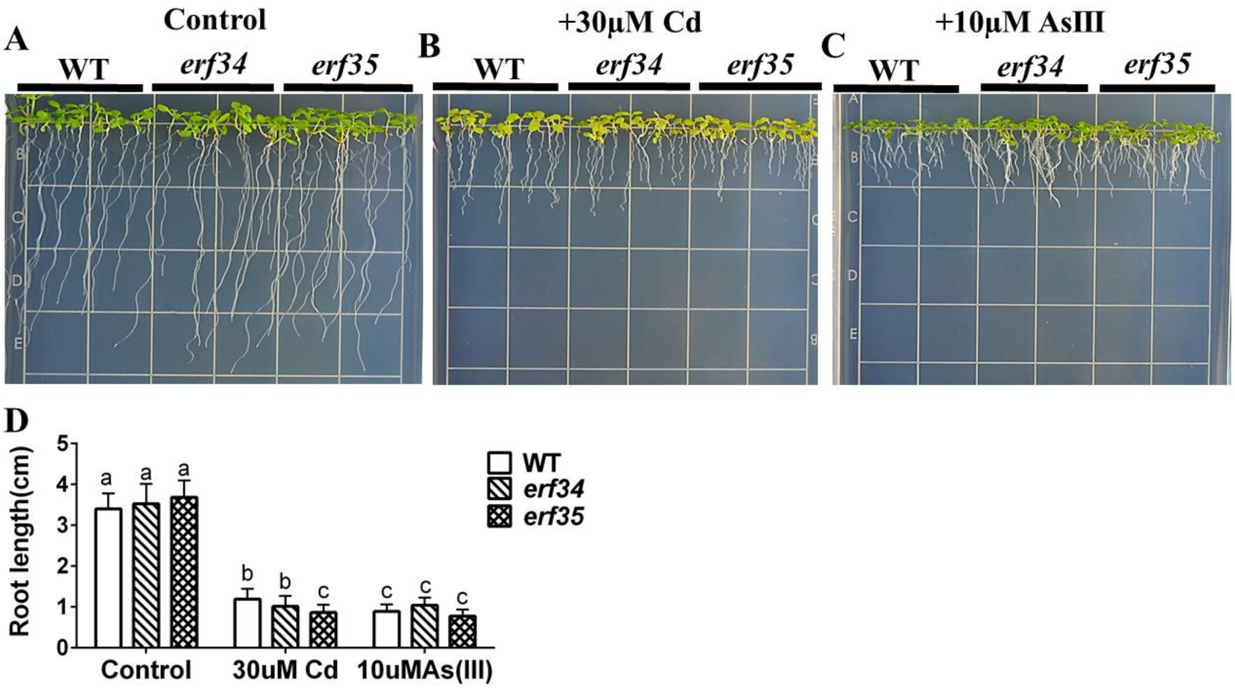


#### Supplemental Figure 3

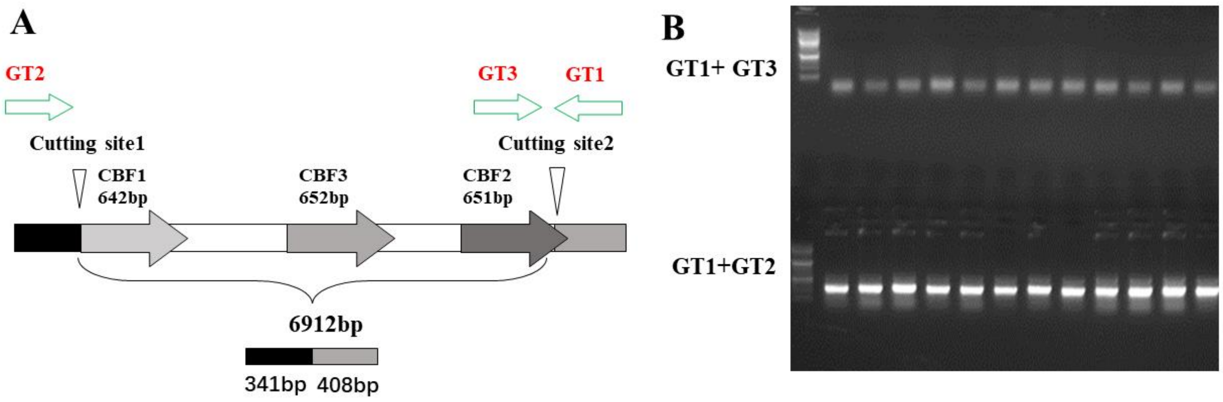

### Supplemental Figure 4

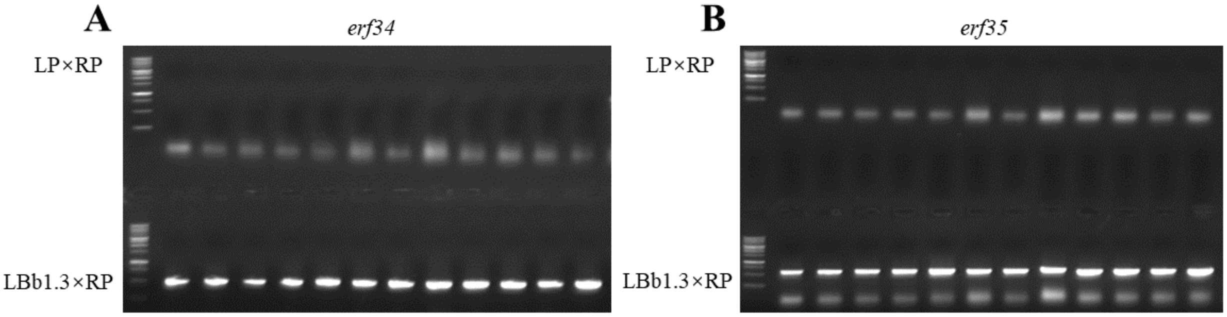


### Supplemental Figure 5

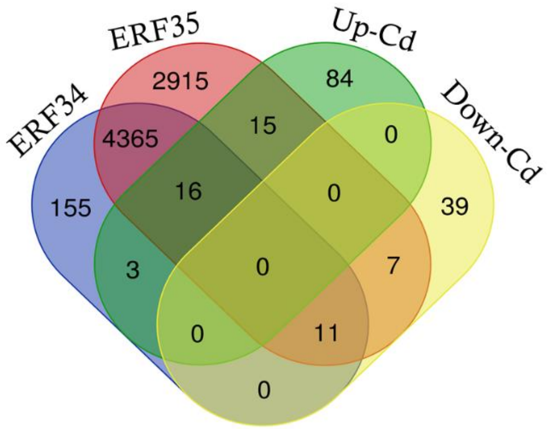

### Supplemental Figure 6

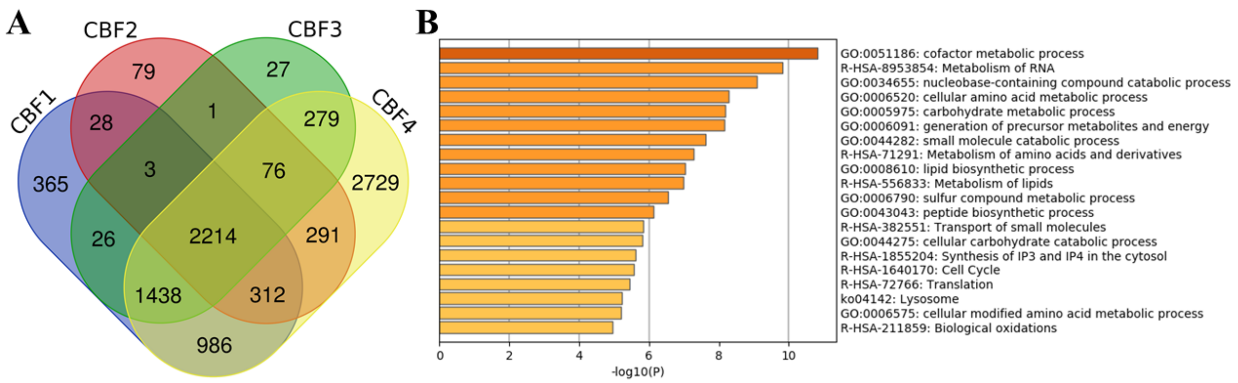
